# Bilayer acoustic force spectroscopy (BAFS) for quantifying receptor-antigen binding strength in immune synapses

**DOI:** 10.64898/2026.03.23.713630

**Authors:** Nebojsa Jukic, Tom M.J. Evers, Anne Walters, Chikim Nguyen, Mai Vuong, A. Christina Heroven, Ricardo A. Fernandes, Sander J. Tans, Kristina A. Ganzinger

## Abstract

Immune cell receptor – ligand interactions are key to cancer immunotherapy. However, receptor-ligand affinities often fail to predict T-cell mediated cancer killing, while immune-target cell binding strength measurements are limited by low precision and high non-specific binding. Here we present bilayer acoustic force spectroscopy (BAFS), a method to quantify the binding strength of receptors in immune synapses that virtually eliminates non-specific binding and increases the resolving power by up to 50-fold. By replacing target cells with a supported lipid bilayer functionalized with antigens, BAFS avoids antigen-independent interactions and target cell heterogeneity, while maintaining the spatial self-organization of receptors that typifies active immune synapses. We demonstrate the high sensitivity and control by showing how CAR T-cell synapse strength depends on CD19 antigen density, and by revealing that CD8 synergistically strengthens αβTCR-pMHC synapses independently of Lck recruitment to CD8. BAFS is a general method that can be used broadly in immunotherapy screening and to dissect the complex molecular interactions that underpin immune synapse activation.

## Introduction

Adoptive cell transfer immunotherapies, especially T cell therapies with modified T cell receptors (TCRs)^1^ or chimeric antigen receptors (CARs)^2^, have transformed the cancer treatment landscape, offering durable responses in patients with otherwise refractory disease^3^. They build on the ability of cytotoxic lymphocytes to selectively recognize tumor-associated antigens and eliminate malignant target cells. However, despite remarkable clinical advances, challenges in optimizing and mechanistically dissecting target cell recognition hamper further progress in developing improved cell-based immunotherapies. Screening pipelines can evaluate thousands of receptor variants for antigen specificity and affinity, but often fail to predict *in vivo* performance of receptor candidates^4^. One reason is that receptor-antigen affinities alone do not capture the spatiotemporal patterns that are thought to underlie successful TCR and CAR-T immune synapse formation, including receptor organization into microclusters and their driven movement to the synapse center^5^. Cell-cell binding strength or avidity measurements have emerged as a key alternative predictor of functional outcome, as recently recognized by the United States Food and Drug Administration^6^. They encompass the spatiotemporal organization of receptor-antigen pairs that underlies functional immune synapses, and avidity measured by acoustic force spectroscopy (AFS)^7,8^ was shown to correlate well with cytotoxicity: CAR-T constructs with higher avidity display stronger cytokine release, tumor lysis, and *in vivo* tumor control, while lower avidity is associated with impaired effector function^9–11^.

However, current AFS implementations measuring cell-cell interactions have drawbacks that severely limit their potential. First, immune-target cell binding involves a highly diverse and complex molecular interaction landscape, which masks the individual immune receptor contributions that are key to immune activation^12,13^. The vast number of interactions, their varying effect on receptor signaling, and the feedback between cell adhesion and receptor activation further exacerbate masking effects^14–17^. For instance, even if cells interact strongly via interactions like LFA-1–ICAM-1 or CD2–CD58, they may lack proper CAR engagement and hence show minimal antigen-specific killing. Finally, target cells can be highly heterogeneous, both between measurements and within the target cell monolayers. Heterogeneity is partly due to variability in adhesion molecule expression levels and integrin activation state, and introduces significant measurement uncertainty^18^. This lack of control and inability to disentangle effects is problematic both for mechanistic studies and immune receptor optimization use cases.

Here, we address these limitations by introducing Bilayer Acoustic Force Spectroscopy (BAFS): a novel, lipid bilayer-based experimental platform for cell avidity measurements. Supported lipid bilayers (SLBs) functionalized with antigens can mimic target cells, and faithfully recapitulate key aspects of immune synapse formation when interacting with immune cells, including receptor organization and activation^19,20^. BAFS provides precise control of target ligand identity, with negligible non-specific interactions, and thus reveals the avidity contributions of receptor-antigen interactions alone. It strongly limits target variability and shows orders of magnitude increases in signal-to-noise ratio (SNR) compared to cell-cell avidity approaches, thus revealing synapse binding strength differences that could not be detected previously. BAFS allows controlled variation of antigen density and can be used generally for any combination of target ligand and cell types, as shown here for CAR and TCR mediated interactions. Illustrating how the control and precision of BAFS can be used to dissect the complexity of immune receptor interactions, we show that the co-stimulatory protein CD8 acts synergistically with αβTCR receptors in binding pMHC ligands, and that it does so independently of CD8 recruitment of Lck. BAFS offers a new platform for screening and mechanistic studies of natural and engineered TCRs and CARs and can be more broadly applied to dissect the contributions of any receptor-ligand pair involved in any immune synapse or cell-cell binding of interest.

### Establishing Bilayer Acoustic Force Spectroscopy (BAFS)

The BAFS method comprises the following steps (**Fig. 1a,b**). An SLB is first formed in a microfluidic AFS chip (z-Movi, LUMICKS, Amsterdam) by flowing in membrane vesicles, which is then functionalized with a ligand of choice, using a low fraction of nickelated lipids in the SLB. Here we used His-tagged CD19 extracellular domain as the paradigmatic CAR T target antigen. Effector cells, here CD19-targeting CAT CAR^21^ expressing Jurkat T cells (CAR+ T cells), are flowed in and incubated with the SLBs. A piezoelectric element within the microfluidic chip is operated near its resonant frequency, causing an acoustic force that drives effector cell detachment from the bilayer in band-like regions (**Fig. 1a**,**c**). Finally, the percentage of cells that remain bound after a force ramp to 1000 pN serves as a quantitative proxy of cellular avidity (**Fig. 1d**). Measurement replicates are obtained using multiple bilayer-functionalized AFS chips run in parallel (as in Fig. 1d), or by reusing the same bilayer-functionalized chip and injecting fresh effector cells between runs – the latter resulting in similar binding curves and endpoint values at 1000 pN, with only slightly increased inter-run variability (**Extended Data Fig. 1, Supplementary Note 1**).

**Fig. 1.**
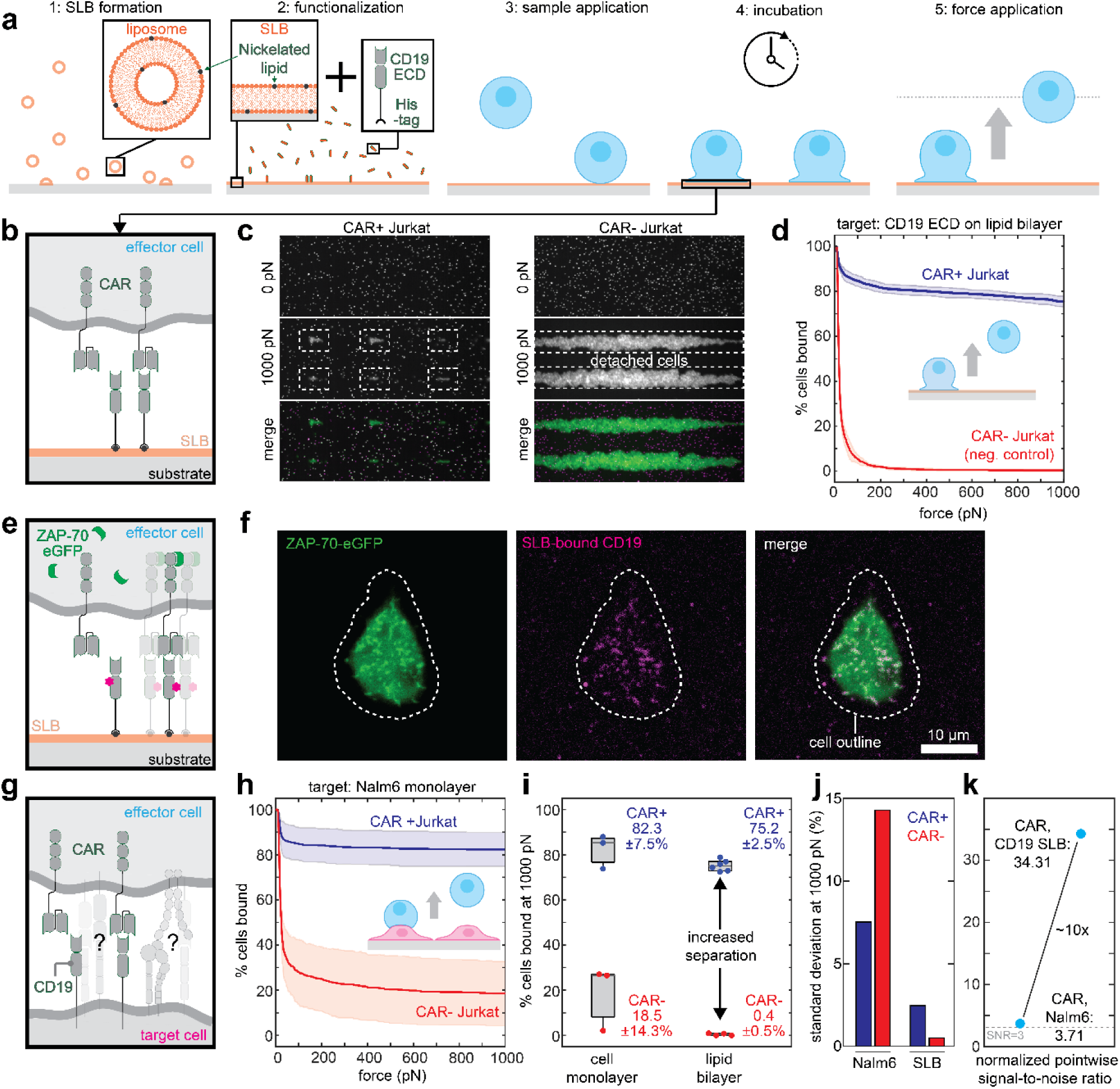
Bilayer Acoustic Force Spectroscopy (BAFS). **a,** BAFS sample preparation: step 1 – supported lipid bilayers are formed by injection of synthetic liposomes into AFS microfluidic chip; step 2 – passivated supported lipid bilayers are functionalized by injection of target ligand into the AFS chip; step 3 – after washing out excess ligand, sample cells (effector or control) are injected into the AFS chip; step 4 – sample cells are incubated on functionalized bilayer before washing out unadhered cells; step 5 – generation of acoustic standing waves, with a force ramp ranging from 0 to 1000 pN, leads to partial detachment of sample cells from the functionalized lipid bilayer. **b,** Schematic of cell attachment to functionalized SLBs due to formation of reconstituted immune synapses specifically through receptor-ligand interaction. **c,** Comparison of sample (CAR+ Jurkat T cells) and control (CAR- Jurkat T cells) at 0 pN (top) and 1000 pN (middle). CAR- Jurkat T cells are almost completely detached at 1000 pN, whereas most CAR+ Jurkat T cells are still attached at 1000 pN. Merge: green – detached cells collecting in acoustic nodes in the chip, magenta – cells attached at 0 pN but not 1000 pN, white – cells attached at 0 pN that remain attached at 1000 pN. **d,** Result of BAFS avidity measurement: average percentage of CAR+ Jurkat T cells (blue, n = 6 runs) and CAR- Jurkat T cells (red, n = 4 runs) bound to CD19+ SLB as a function of force. Shaded area: standard deviation. **e,** Schematic of Jurkat T cells expressing ZAP-70-eGFP in contact with SLBs functionalized with fluorescently labelled CD19. **f,** Example confocal microscopy images showing a CAR+ Jurkat T cell expressing ZAP-70-eGFP bound to CD19+ SLB after force application. ZAP-70-eGFP clusters in the effector cell (green, left) and SLB-bound CD19 clusters (magenta, middle) spatially colocalize (merge, right), indicating that increased binding is due to CD19-CAR interaction. **g,** Schematic of topographically complex cell-cell contact sites comprising not only specific receptor-ligand interactions, but also interactions between adhesion molecules and other off-target interactions. **h,** Result of cell-cell avidity measurement (classic AFS): average percentage of CAR+ Jurkat T cells (blue, n = 3 runs) and CAR- Jurkat T cells (red, n = 3 runs) bound to CD19-expressing Nalm-6 cells as a function of force. Shaded area: standard deviation. **i,** Boxplot comparing CAR+ Jurkat T-cell binding (blue, left) and CAR- Jurkat T-cell binding (red, left) to Nalm-6 cells at 1000 pN to CAR+ Jurkat T-cell binding (blue, right) and CAR- Jurkat T-cell binding (red, right) to CD19+ SLB at 1000 pN. **j,** Standard deviation of average binding at 1000 pN is decreased about 3-fold for CAR+ Jurkat T cells (blue) in BAFS as compared to AFS, and ∼29-fold for CAR- Jurkat T cells (red), due to abolished non-specific binding on the CD19+ SLB. **k,** Normalized pointwise signal-to-noise ratio is increased ∼10-fold in BAFS (SNR = 34.31) as compared to AFS (3.71). Dotted line: SNR=3 threshold (approximately 99.7% confidence that a detected signal is real and not a random fluctuation of background noise).

CAR+ T cells were found to strongly bind the target SLBs, with over 75.2 ± 2.5% (mean and SD) of the cells remaining bound at 1000 pN (**Fig. 1d**, blue). Notably, binding of control Jurkat cells not expressing CARs (CAR- T cells) was negligible, with 0.4 ± 0.5% of the cells remaining bound at 1000 pN (**Fig. 1d**, red). Thus, the CAR+ T cell data quantified avidity solely driven by CAR expression and CAR binding to its cognate ligand. Confocal microscopy performed after these measurements confirmed key features of immune synapse formation. Specifically, when fluorescently labelled ZAP-70, a tyrosine kinase that binds phosphorylated CARs, was expressed in CAR+ T cells (**Fig. 1e**), we observed ZAP-70 microclusters (**Fig. 1f**, left) colocalizing with microclusters of fluorescently labelled CD19 on the lipid bilayer (**Fig. 1f**, middle). This key step in T-cell activation is a characteristic sign of immune synapse formation (**Fig. 1f**, right). Lipids in the SLBs were found to be mobile, using fluorescence recovery after photobleaching assays in a custom-built total internal reflected fluorescence (TIRF) microscope^22^ (**Extended Data Fig. 2**). SLB mobility is consistent with the need for ligand diffusion for proper synapse formation^23^. We recapitulated these results by measuring CAR+ and CAR- binding on SLBs through biotin moieties rather than nickelated lipids (**Extended Data Fig. 3**). This confirmed that BAFS is compatible with multiple surface functionalization chemistries.

### BAFS improves resolution and decreases variability in avidity measurements

Next, we tested whether BAFS improves upon cell-based AFS using the same effector cells (**Fig. 1a-d**). In the latter, the AFS chip glass surface is functionalized with a target cell monolayer^24^, for which we use human B cell precursor leukemia Nalm-6 cells that natively express CD19 (**Fig. 1g**). We found that 82.3 ± 7.5% (mean and SD) CAR+ T cells remained bound at 1000 pN, compared to 18.5 ± 14.3% for the negative control (**Fig. 1h**). Hence, BAFS showed similar mean endpoint values for CAR+ cells as compared to AFS but yielded a 46-fold lower non-specific binding by CAR- cells, and hence increased separation between control and sample (**Fig. 1i**, red; **Extended Data Table 1; Supplementary Movie 1**). In addition, BAFS showed large decreases in variation between measurements; 29-fold for CAR- cells (14.3% to 0.5%) and a 3-fold decrease for CAR+ T cells (7.5% to 2.5%, **Fig. 1j**). Together, these data indicated a signal-to-noise ratio (SNR) of ∼34 for BAFS compared to ∼4 for AFS (**Fig. 1k**). Together, our findings thus far show that BAFS allows us to quantify the avidity of clinically used immunoreceptors, such as CARs, at superior resolution and minimal background compared to AFS based on cell monolayers.

The high SNR raised the question if BAFS could detect much smaller cell adhesion differences. Specifically, we contrasted a bilayer with nickelated lipids only, which are proposed to cause weak (electrostatic) binding of Jurkat T cells^25^, and a bilayer where this effect is masked by binding His-tagged SNAP-tag molecules to the nickel headgroups (**Fig. 2a**). Both showed a negligible fraction of CAR+ T cells that remained bound at 1000 pN, with no cells remaining bound on the masked SLB (**Fig. 2a**, yellow) and 2.8 ± 1.4% bound on the unmasked SLB (**Fig. 2a**, gray). However, the binding curves displayed a distinct divergence below 500 pN, with more cells remaining bound to the unmasked SLB at these low forces. The force at which 90% of the cells were unbound increased 4-fold (from 40 pN to 179 pN, **Fig. 2a**, inset, dot) in the unmasked SLB compared to the masked SLB, even as the median unbinding force remained similar (16 pN vs. 24 pN, **Fig. 2a**, inset, square). These data showed that BAFS can exploit the low-force regime that is typically not used in AFS due to insufficient resolution and lack of reproducibility. Thus, BAFS provides a sensitive tool to dissect low-force contributions to cell-cell adhesion.

**Fig. 2.**
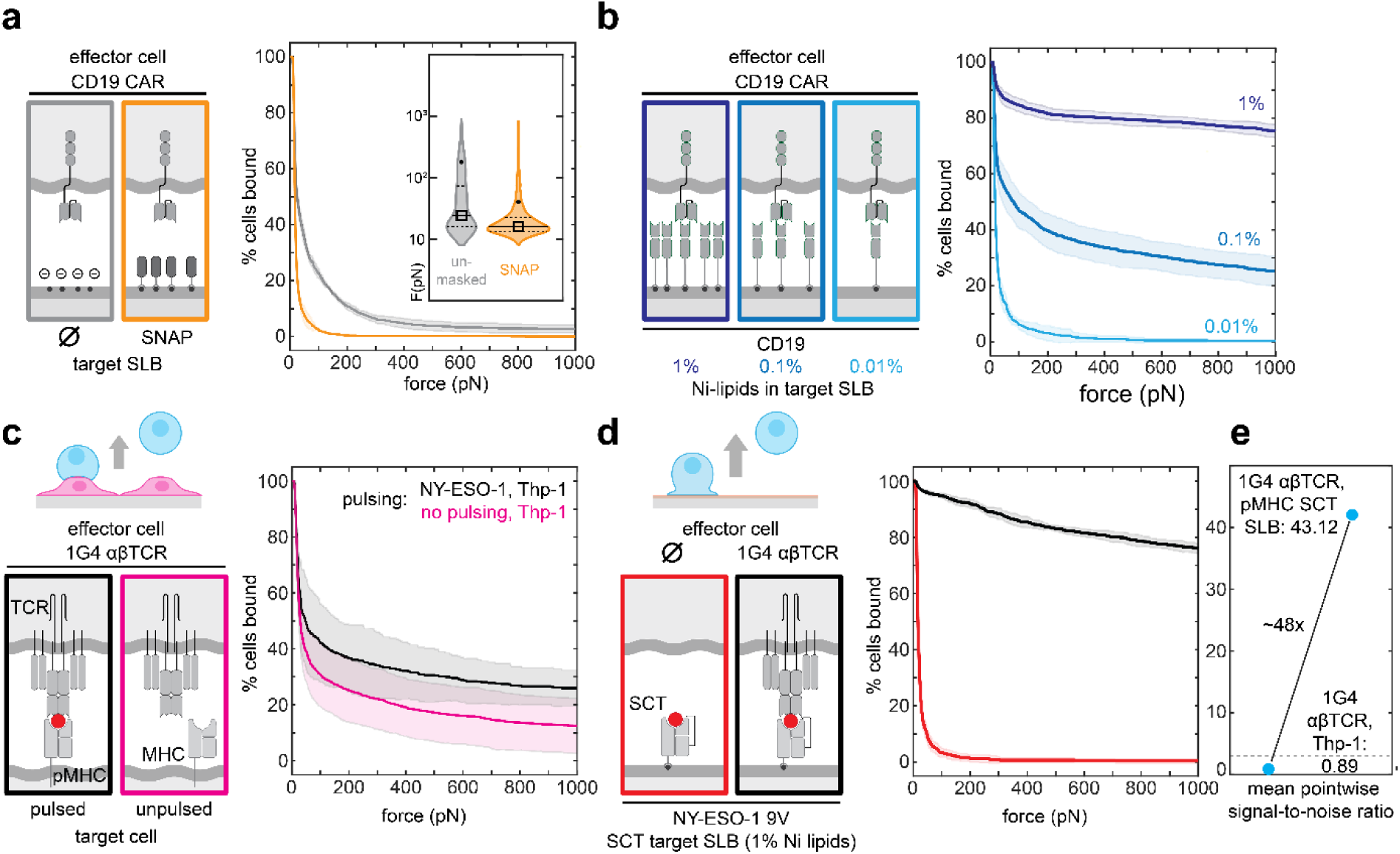
BAFS provides ligand control and can probe diverse immune interactions. **a,** Unmasked nickelated lipids induce weak, non-specific binding of CAR+ Jurkat T cells. Right: Average percentage of CAR+ Jurkat T cells bound to CD19- SLB (gray, n = 4 runs) or SNAP-tag+ SLB (yellow, n = 2 runs) as a function of force. Shaded area: standard deviation. Inset: violin plots showing rupture force distribution for all cells tracked across the z-Movi BAFS measurements (gray: 1609 cells tracked in 4 separate runs, yellow: 1227 cells tracked in 2 separate runs); square – median unbinding force; dot – 90% unbinding force. **b,** Titration allows insights into ligand concentration-dependent binding effects. Average binding curves (mean±SD) of CAR+ Jurkat cells as a function of force on SLBs with 1% (dark blue, n = 6 runs), 0.1% (blue, n = 4 runs) and 0.01% (cyan, n = 3 runs) nickelated lipids, functionalized using the same concentration of His-tagged CD19 ECD. **c,** Average binding curves (mean±SD) of 1G4 αβTCR+ Jurkat T-cells on target Thp-1 cells after pulsing with cognate NY-ESO-1 antigen (black, n = 4 runs) and untreated Thp-1 cells (magenta, n = 4 runs). Deriving conclusion based on statistical analysis is rendered difficult due to the weak separation of the curves and large SD (*P*=0.062, two-sample t-test, at 1000 pN). For detailed statistical analysis, see **Extended Data Figure 4. d,** Average binding curves (mean±SD) of 1G4 αβTCR+ Jurkat T-cells on NY-ESO-1 SCT+ SLBs (black, n = 3 runs) and αβTCR-Jurkat T-cells on NY-ESO-1 SCT+ SLBs (red, n = 4 runs). The separation at 1000 pN and small variation between runs for each condition permits clear separation of sample and control. **e,** Mean pointwise signal-to-noise ratio is increased ∼48-fold in BAFS (SNR = 43.12) as compared to AFS (0.89). Dotted line: SNR=3 threshold.

### BAFS allows control of ligand surface density

An advantage of BAFS is the ability to precisely control ligand surface density, which is challenging to achieve when using cell monolayers as binding targets. We varied the density of CD19 by decreasing the nickelated lipid abundance over two orders of magnitude (from 1% to 0.01%) while keeping the concentration of the injected (His-tagged) CD19 constant. The percentage of cells bound at 1000 pN dropped from 75.2 ± 2.5% (1% Ni lipids, **Fig. 2b**, dark blue) to 25.3 ± 5.2% (0.1% Ni lipids, **Fig. 2b**, blue), while binding was almost completely abolished on an SLB containing 0.01% nickelated lipids (0.29 ± 0.5%, **Fig. 2b**, cyan). Overall, these experiments illustrate that BAFS provides the measurement sensitivity and experimental control to titrate antigen density, and shows that immune synapse binding strength depends strongly on the antigen surface density.

### BAFS can probe diverse receptor-ligand interactions central to immune synapses

To test BAFS generality and resolving power, we turned to studying αβTCR binding. First, we used cell-based AFS to obtain a baseline. A target cell monolayer of human leukemia monocytic cell line Thp-1 was formed in an AFS chip. These cells bind and activate 1G4 αβTCR+ Jurkat T cells (TCR+ T cells), when incubated (pulsed) with the NY-ESO-1 peptide^26^. The percentage of TCR+ cells bound to the target cells at 1000 pN was 25.9 ± 6.4% (**Fig. 2c**, black). Notably, this value did not differ significantly from control data on a target cell monolayer not treated with the cognate antigen peptide (12.5 ± 9.8%; **Fig. 2c**, magenta; see **Extended Data Fig. 4a,b** for detailed statistical analysis), showing that this TCR-specific binding is challenging to resolve with cell-based AFS. When substituting the target monolayer cell line Thp-1 for NY-ESO-1 pulsed Nalm-6 cells, the percentage of TCR+ T cells bound increased to 52.5 ± 3.4%, while the unpulsed control also increased substantially (25.4 ± 7.7%; **Extended Data Fig. 4c**), highlighting the variability of background binding and in general the large influence of target cell lines on cell-cell avidity measurements.

Next, we performed BAFS measurements using SLBs functionalized with the cognate pMHC (NY-ESO-1 9V) as single chain trimer (SCT). Here, the percentage of TCR+ cells bound to the SLB at 1000 pN was higher at 75.95 ± 1.75%. Binding was negligible for TCR- T cells, with 0.4 ± 0.5% cells still bound at 1000 pN (**Fig. 2d**), showing again the high resolving power of BAFS. The SNR thus increased dramatically from 0.89 for AFS to 43.12 for BAFS – a 48-fold improvement (**Fig. 2e**). Notably, BAFS also outperformed the target-optimized cell-cell avidity AFS assays in terms of sample-control separation and variance between runs (compare **Fig. 2d**, **Extended Data Fig. 4b-d**, **Extended Data Table 2**). Unlike AFS, BAFS was thus able to specifically quantify the avidity of TCR and cognate pMHC.

### BAFS can dissect complex interactions in immune synapses

We next applied BAFS to address a long-standing question in immunology, namely how the co-receptor CD8 contributes to TCR-pMHC synapse stabilisation. CD8 binds to the α3 domain of MHC class I, distinct from the TCR-binding site of the pMHC^27,28^, which is thought to stabilise short-lived interactions between TCR and cognate pMHC^29–31^. Furthermore, the cytoplasmic domain of CD8 also binds the lymphocyte-specific protein tyrosine kinase (Lck), which phosphorylates and hence binds the TCR/CD3 immunoreceptor tyrosine-based activation motifs on the intracellular side^32^. However, it remains poorly understood how these two mechanisms in this three-way interaction contribute to synapse strength, and if both are needed for CD8 to “clamp” the TCR-pMHC interactions^33–35^.

We first compared 1G4 TCR+ Jurkat T cells that expressed wild-type CD8 (CD8αβ+) to cells that did not (CD8-). We found that both cells similarly showed strong binding (**Fig. 3a,b**, 1% nickelated lipids, orange and black solid lines, respectively). Replacing wild-type CD8 with a CD8 mutant that does not recruit Lck (CD8α C194S/C196S / CD8β WT)^36,37^ did not affect unbinding throughout the force ramp (**Fig. 3a,b**, cyan solid line). Hence, at high pMHC densities, neither CD8 presence nor Lck-CD8 binding contributed to synapse strength, or did so in limited manner. Intriguingly however, cells expressing wild-type CD8 but no TCR (**Fig. 3a,b**, magenta solid line) showed similarly high degrees of binding as the TCR+ cells with or without CD8. This suggested that direct CD8-pMHC interactions do form, and can even be sufficient to generate long-lived binding, despite the low pMHC affinity of CD8 compared to αβTCR (∼200 μM vs ∼7 μM)^29,38,39^. It also raised the idea that in the TCR+ experiments, CD8 did have an effect, but that it was masked by dominant TCR-pMHC interactions, possibly caused by high pMHC densities.

**Fig. 3.**
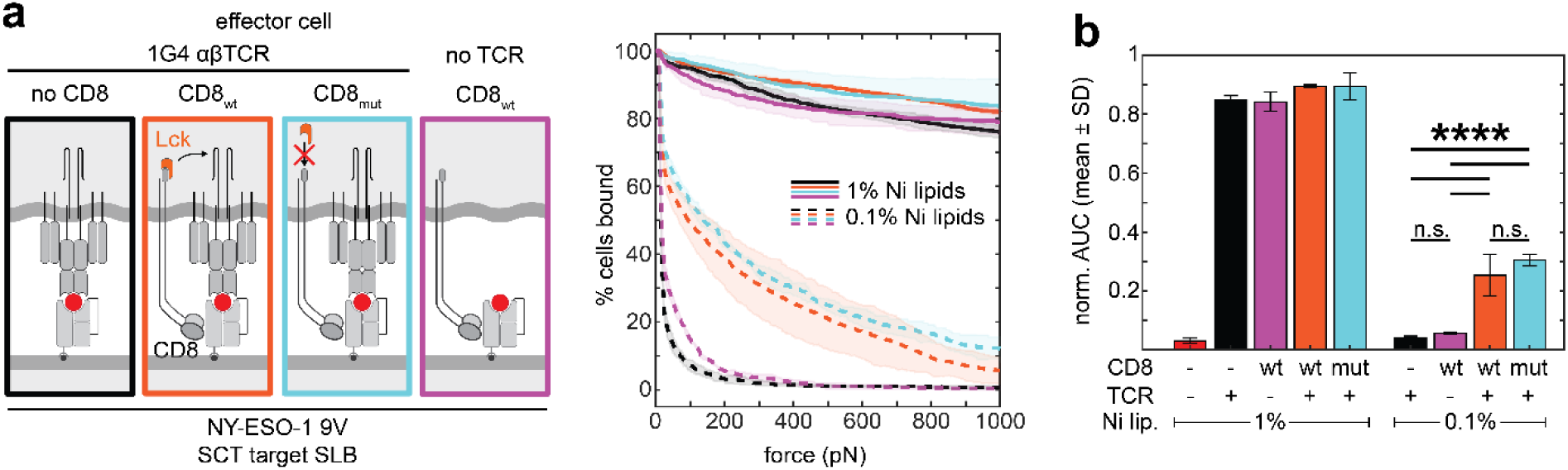
BAFS dissects contributions of CD8 and 1G4 αβTCR on T cell avidity. **a,** Left: Schematic illustrating different avidity measurement conditions, with effector cells expressing 1G4 αβTCR only (black), CD8 wild-type only (magenta), or co-expressing 1G4 αβTCR with either CD8 wild-type (orange) or CD8 mutant deficient in Lck binding (cyan). Right: avidity measurements on 1% (solid lines) and 0.1% (dashed lines) nickelated lipid bilayers. Curves are average values across at least 3 measurements per condition; shaded areas are standard deviation. **b,** Normalized area under the average binding curves for all conditions shown in Fig. 3a, right. Error bars: standard deviation. All conditions n = 3 runs, except TCR-/CD8- (red, identical to Fig. 2d, n = 4 runs). The reported *P*-values were obtained by running a non-parametric log-rank test on Kaplan-Meier survival plots of all individual cells in each replicate, followed by correction for multiple comparisons. *P*-values were reported as significant only if both statistical significance and a threshold criterion for effect size were met; a more detailed analysis is provided in **Extended Data Figure 5.**

Indeed, at a tenfold lower pMHC density (0.1% nickelated lipids), CD8 did increase the binding strength when expressed with TCR, compared to TCR alone (**Fig. 3a**, dashed orange and black, respectively). The difference was most prominent in the low-force regime, with TCR+/CD8+ cells detaching slower than TCR^+^/CD8- cells during the force ramp (**Fig. 3a,b**). Interestingly, cells expressing only CD8 or only TCRs both showed negligible binding (**Fig. 3a,b**, dashed magenta and black), at levels comparable to TCR-/CD8- cells (**Fig. 2d** and **3b**, red). These data thus showed that at low pMHC densities, CD8 and the TCR contribute synergistically (rather than additively) to the synapse binding strength. We also found that the CD8 mutation did not significantly affect binding in TCR+ cells (**Fig. 3a,b**, dashed orange and cyan; see **Extended Data Figure 5** for statistical analysis), indicating that Lck recruitment by CD8 is more important for downstream signaling than for synapse maturation and binding strength. Overall, these experiments highlighted how BAFS can dissect complex contributions to immune synapse formation, which are challenging to disentangle using existing methods.

## Discussion

In summary, we present the BAFS method for quantifying immune synapse binding strength or avidity, which exploits the binding of immune cells to functionalized supported lipid bilayers (SLBs). BAFS allows precise control of ligand identity and density, elimination of antigen-independent interactions, and homogeneous ligand distribution, while overall producing highly reproducible data with minimal need for optimization. It overcomes several drawbacks of existing cell-cell avidity measurement methods. First, it eliminates masking effects from the many other molecular interactions between cells, which may be specific or non-specific, and are only partially known. Hence, it reveals the avidity driven by the receptor-ligand pair(s) of interest, while preserving their spatial organization. Second, it dramatically increases signal-to-noise ratio, which provides unprecedented resolving power at high forces, and opens up previously inaccessible low-force regimes. Third, it offers a level of control and precision that can be used to disentangle the individual contributions of complex interactions between multiple molecules that typify immune synapses, by introducing interactors in a titratable and combinatorial manner.

We illustrated the capabilities of BAFS by quantifying how CAR T-cell immune synapse binding strength depends on the CD19 antigen surface density, by resolving specific binding in αβTCR-pMHC immune synapses, and by dissecting the three-way interplay between αβTCR, pMHC and CD8 in the latter. Although CD8 and TCR 3D affinities for pMHC – as measured by other methods – differ by two orders of magnitude, BAFS can distinguish individual contributions and synergistic effects on synapse strength, thus addressing a fundamental open question in immune cell biology.

BAFS is a general method that can be applied in a wide spectrum of cases. For instance, it can be used to reveal how immune synapse strength and activation is controlled by two or more ligands, using orthogonal functionalization via nickelated and biotinylated lipids. Examples include the LFA-1-ICAM-1 interaction or CD28-CD80/86 costimulatory signals, which T cells exploit besides the TCR-pMHC signal to produce a full effector response^40^, and designer receptors that implement logical gates for CAR activation^41^. Another application area are T-cell engagers (TCEs), which are engineered to increase immune response by binding partners both on the T-cell side (e.g. CD3 in the TCR complex) as well as the target cell side (e.g. CD19), including 2+1 format TCEs that bind two target ligands with reduced affinity for each individual ligand^42^. Furthermore, BAFS can be used to study surface ligand density-dependent phenomena that limit the utility of CAR-T cells in immunotherapy: first, the effect of antigen escape^43^, a process where tumor cells downregulate specific surface antigens to avoid detection and elimination by CAR-T cells, and second, on-target-off-tumor toxicity – a serious side effect during application of high-affinity CARs that detect and are activated by tumor-associated antigens at low densities^44^. Finally, by eliminating the high non-specific background binding and the complex multivalent interactions present at cell-cell contacts, BAFS can enable early separation and elimination of true non-binders from low-avidity binders in panels of candidate constructs during immunotherapy development, which can otherwise be difficult to distinguish. More broadly, BAFS may be used for any cell-cell interactions, ranging from B-cell, NK-cell, and gamma-delta T-cell receptors, to adhesion and co-stimulatory proteins, as well as to immune checkpoint inhibitors.

While ligand control is an important advantage of BAFS, it also limits its application to interactions for which the ligands are known and can be purified for SLB functionalization. These ligands often comprise only the extracellular domains of the actual target proteins, missing transmembrane and cytosolic domains that may allow for ligand conformational changes, oligomerization, or interaction with cytoskeletal components in target cells, and are potentially important to some receptor-ligand interactions *in vivo*^45^. Polymer-cushioned supported lipid membranes may be used to overcome this limitation^46^.

BAFS may also be combined with high-resolution microscopy for observations of reconstituted immune synapses, as illustrated in **Fig. 1**. While correlated force spectroscopy and fluorescence imaging using immersion objectives is not possible yet due to force dissipation when the chip is in contact with immersion liquid^8^, simultaneous force spectroscopy and confocal microscopy would enable more detailed correlation between synapse strength, structure, and immune activation, either statically or as a function of time.

To conclude, BAFS represents an experimental platform that expands the parameter space that can be explored during avidity measurements and greatly increases reproducibility and sensitivity of synapse binding strength measurements. These capabilities are of importance both to users in translational research focusing on resolving power and reproducibility when screening immunotherapy candidates and in quality control maintenance, as well as to users exploring more fundamental aspects of effector cells interacting with their ligands on the target cell surface.

## Extended Data

**Extended Data Fig. 1.**
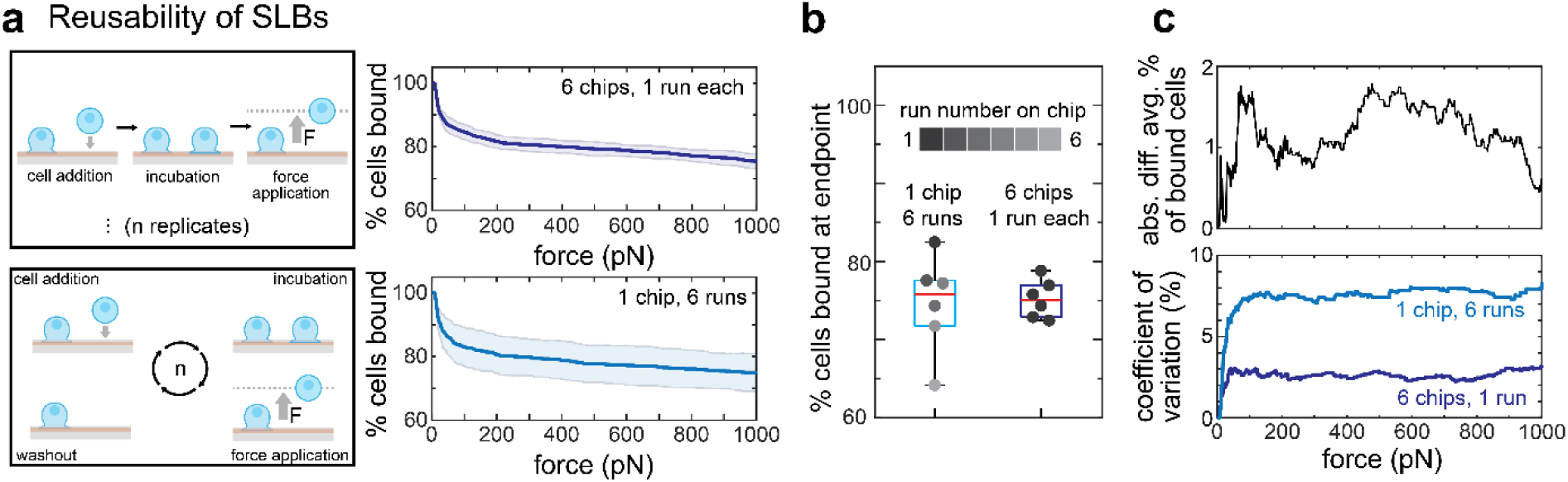
Reusability of functionalized SLBs. **a,** Schematic demonstrating the sample preparation of parallel avidity measurements on several chips functionalized using the same protocol (top left) as compared to sequential avidity measurements on the same SLB-functionalized chip (bottom left). The corresponding average percentage of CAR+ Jurkat T cells bound to a 1% nickelated CD19+ SLB as a function of force (mean±SD) is displayed on the top right (dark blue) and bottom right (light blue), respectively. **b,** Boxplots displaying spread of percentage of cells bound at 1000 pN for sequential (left, light blue) and parallel avidity measurements (right, dark blue), corresponding to runs summarized in the average binding curves in Extended Data Fig. 2a, right. Box: 2^nd^ and 3^rd^ quartile, whiskers: 1^st^ and 4^th^ quartile. Red line: median. Data points in overlaid scatterplot are colored by run order, from dark to light gray. **c,** Top: Absolute difference of average percentage of cells bound as a function of force between sequential and parallel avidity runs. Bottom: coefficient of variation for parallel (dark blue) and sequential (light blue) BAFS avidity measurements, corresponding to the data displayed in the average binding curves in Extended Data Fig. 1a, right.

**Extended Data Fig. 2.**
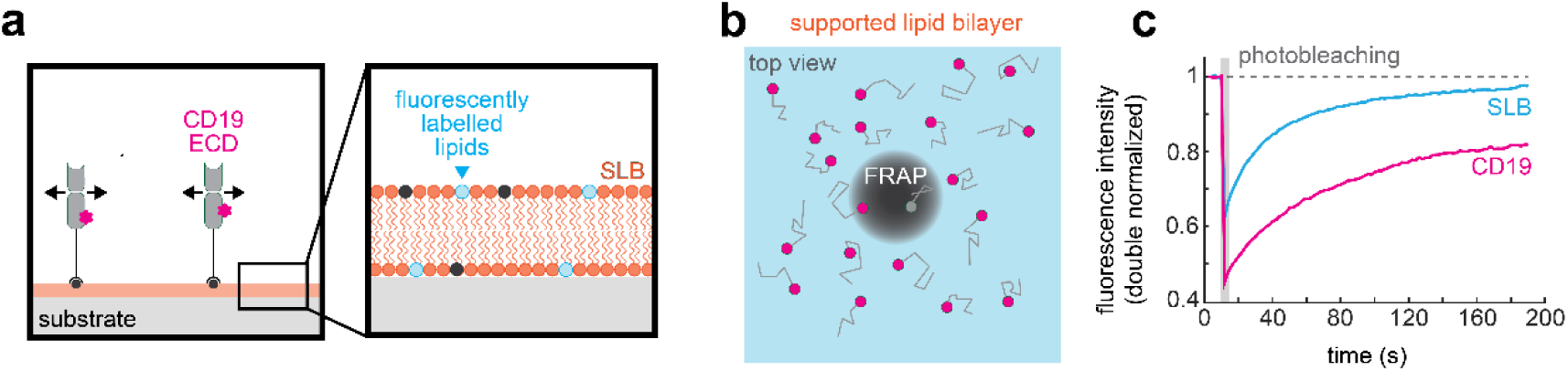
Lipid diffusivity in SLBs. Fluorescence recovery after photobleaching (FRAP) of fluorescently labeled CD19 bound to a fluorescently labeled, partially nickelated SLB confirms bilayer and ligand mobility. **a,** Schematic illustrating the expected lateral diffusivity of CD19 ECD on the SLB, and the composition of the partially nickelated and partially fluorescently labelled SLB. **b,** Schematic illustrating a bleached spot in the SLB, as observed by total internal reflection fluorescence (TIRF) microscopy, and trajectories depicting diffusion of unbleached CD19 ECD into the bleached area. **c,** Quantification of FRAP of fluorescently labeled phospholipids in the SLB (cyan) and the His-tagged fluorescently labeled CD19 functionalized on the SLB (magenta) as a function of time. Intensity curves were double-normalized to an unbleached area of the SLB outside of the FRAP area.

**Extended Data Fig. 3.**
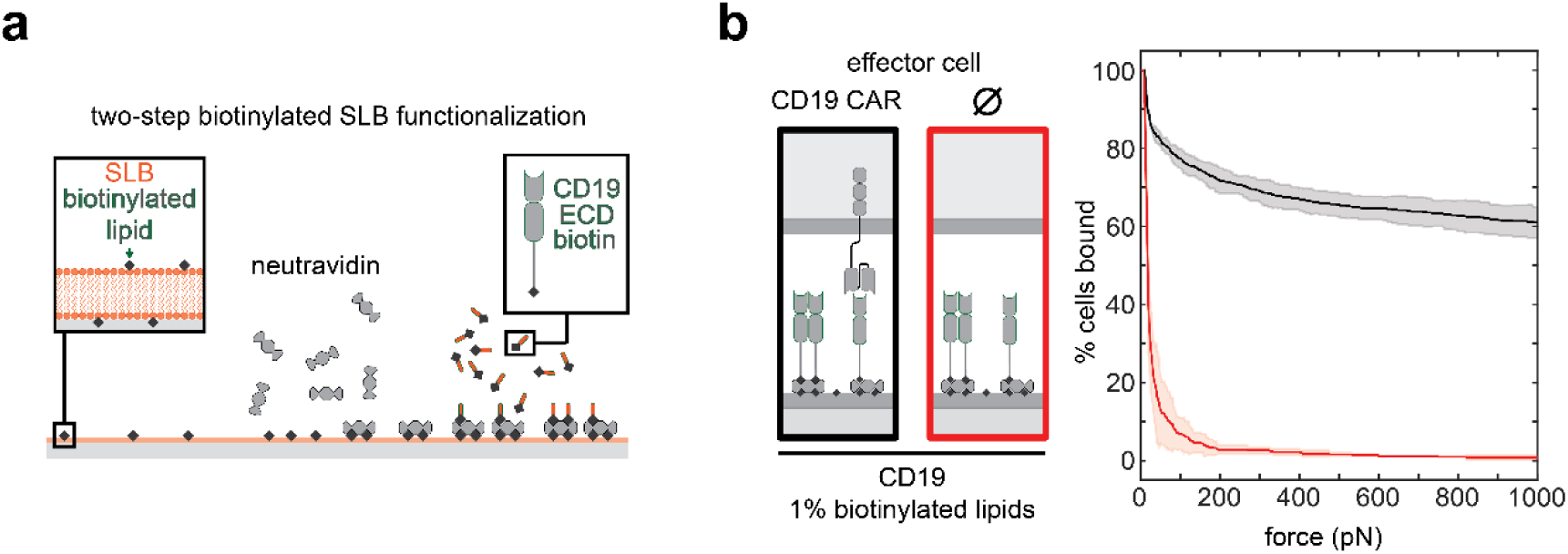
BAFS is compatible with multiple SLB functionalization chemistries. **a,** Schematic illustrating two-step functionalization of 1% biotinylated lipid-containing SLB with CD19 ECD. The lipid bilayer is first functionalized with neutravidin, which binds to the biotinylated phospholipid headgroups on the bilayer surface. In a second step, biotinylated CD19 ECD is flushed into the microfluidic AFS chip. Further sample preparation and avidity measurements are identical to those described in Fig. 1a (see also Methods section). **b,** CAR+ and CAR- cell binding measured on 1% biotinylated SLBs displaying CD19 ECD. Average binding (solid lines) and standard deviation (shaded areas): CAR+ (black, n = 3 runs, sequential measurements) 61.0±4.0%, CAR- (red, n = 2 runs, sequential measurements) 0.8±0.6%.

**Extended Data Fig. 4.**
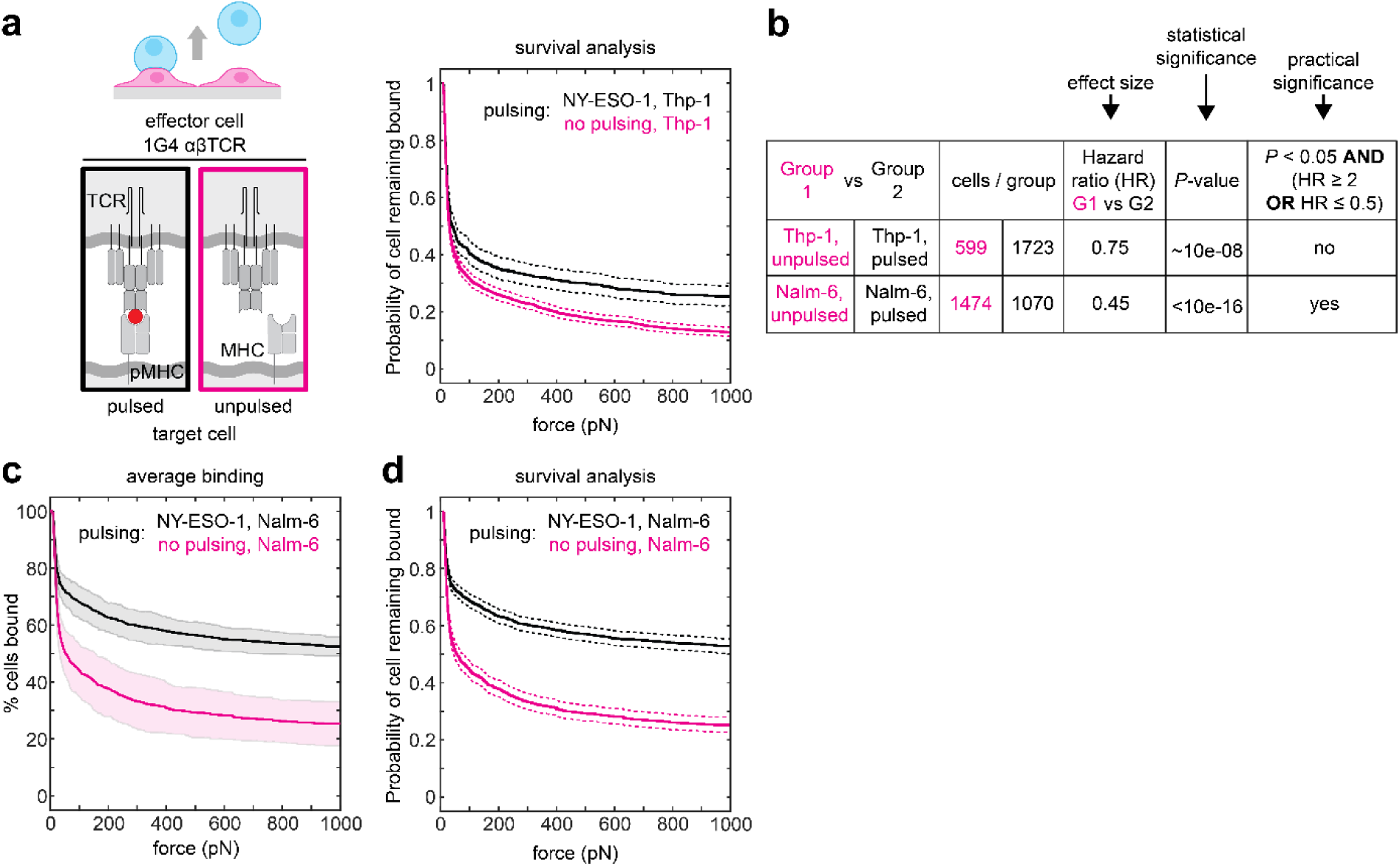
Separation of avidity measurements using AFS can be improved through optimization of assay conditions. **a**, Kaplan-Meier (KM) survival plot of 1G4 αβTCR+ Jurkat cells unbinding from a target cell monolayer consisting of Thp-1 cells that were either pulsed with NY-ESO-1 9V (black) or left untreated (magenta) prior to coincubation of effector cells. Dotted lines: 95% confidence intervals estimated using Greenwood’s formula. **b,** Table displaying statistical analysis using a non-parametric log-rank test. Statistical significance is contextualized using the hazard ratio (HR) as a proxy for effect size; survival curve differences were deemed practically significant only if *P*-value < 0.05 and HR either ≥ 2 or ≤ 0.5. Statistical analysis of survival curves displayed both for effector cells incubated on Thp-1 (top row) or Nalm-6 (bottom row) target monolayers using the same pulsing protocol. **c,** Average binding curves (mean±SD) of 1G4 αβTCR+ Jurkat T-cells on target Nalm-6 cells after pulsing with cognate NY-ESO-1 antigen for 16 h (black) and untreated Nalm-6 cells (magenta). Replacing Thp-1 as the target cell line with Nalm-6 leads to better separation, but background binding of 1G4 αβTCR+ Jurkat T-cells as well as inter-run variability remained high, decreasing the statistical power of the findings. For detailed performance metrics on this dataset and comparison to a BAFS-based approach, see **Extended Data Table 2**. **d,** survival plots for data shown in Extended Data Fig. 4c. Dotted lines: 95% confidence intervals estimated using Greenwood’s formula.

**Extended Data Fig. 5.**
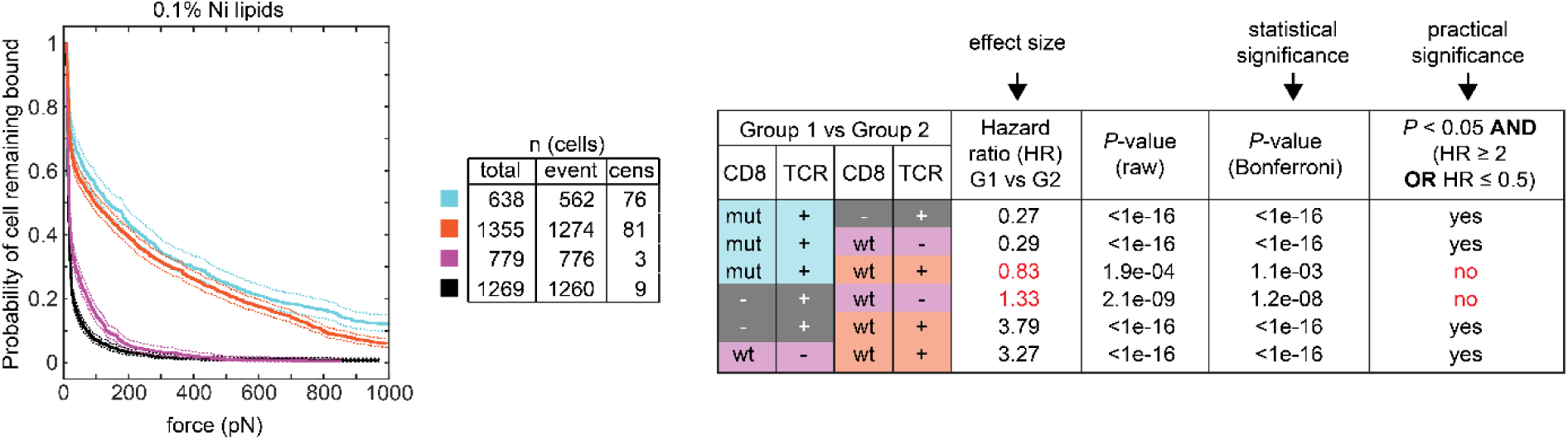
Survival analysis of Jurkat T cells expressing CD8, 1G4 αβTCR, or both, unbinding from bilayers as measured by BAFS. Left: Kaplan-Meier (KM) survival plot of CD8-/TCR+ (black), CD8wt+/TCR- (magenta), CD8wt+/TCR+ (yellow) and CD8mut+/TCR+ (cyan) Jurkat T cells upon unbinding from a 0.1% nickelated bilayer functionalized with NY-ESO-1 SCT. Dotted lines: 95% confidence intervals estimated using Greenwood’s formula. Middle: Table providing total number of cells analyzed per condition (each condition n = 3 replicates), number of cells that detached from the bilayer during force application (event), as well as number of cells remaining bound at 1000 pN (censored). Right: Table displaying statistical analysis using a non-parametric log-rank test. Pairwise survival curve differences were deemed practically significant only if Bonferroni-corrected *P*-value < 0.05 and effect size criterion (HR) either ≥ 2 or ≤ 0.5.

**Extended Data Table 1:**
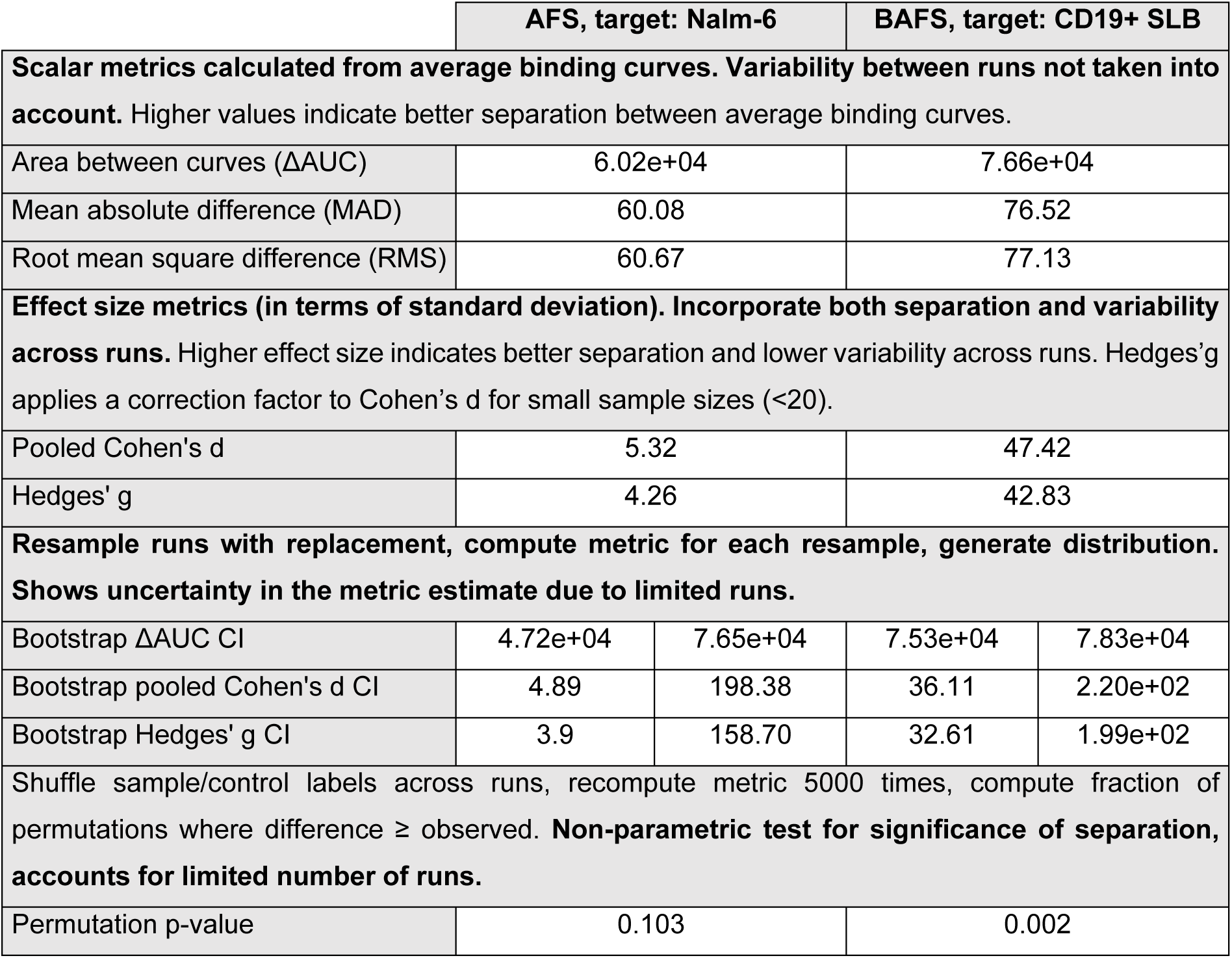
Comparison of performance metrics for CAT CAR+ cell binding measurements using AFS and BAFS. AFS: CAT CAR+ cells binding to Nalm-6 target cell monolayer (see Fig. 1g,h). BAFS: CAT CAR+ cells binding to His-tag CD19 ECD functionalized 1% nickelated bilayer (see Fig. 1d). The two experimental approaches were compared for scalar metrics (area between curves – ΔAUC, mean absolute difference – MAD and root mean square difference – RMS), with a larger difference indicating better separation between average binding curves across the methods. Comparison of effect size metrics (Pooled Cohen’s d and Hedges’ g) allows interpretation of both separation as well as variability across runs, with higher values indicating better separation between average binding curves and lower variability between runs. By convention, Cohen’s d ≤ 0.2 small effect size, ≤ 0.5 medium effect size, ≤ 0.8 large effect size; both for AFS as well as BAFS the effect size is extremely large. Bootstrapping approaches were used to calculate the uncertainty in the metric estimate due to the limited number of runs on each chip. Finally, a non-parametric test was used to ascertain the significance of the separation in AFS and BAFS, showing a *P*-value of 0.1027 for AFS (not statistically significant) vs 0.0017 for BAFS.

**Extended Data Table 2:** Comparison of performance metrics for 1G4 αβTCR+ cell binding measurements using unoptimized AFS, target-optimized AFS and out-of-the box BAFS. AFS: TCR^+^/CD8^-^ cells binding to NY-ESO-1 9V pulsed Thp-1 (unoptimized, left column; see Fig. 2c) or Nalm-6 (optimized, middle column; see Extended Data Fig. 4) target cell monolayer. BAFS: TCR^+^/CD8^-^ cells binding to HLA-A02:01 NY-ESO-1 9V SCT functionalized 1% nickelated bilayer (see Fig. 2d). The two experimental approaches were compared for scalar metrics (area between curves – ΔAUC, mean absolute difference – MAD and root mean square difference – RMSD), with a larger difference indicating better separation between average binding curves across the methods. Comparison of effect size metrics (Pooled Cohen’s d and Hedges’ g) allows interpretation of both separation as well as variability across runs, with higher values indicating better separation between average binding curves and lower variability between runs. By convention, Cohen’s d ≤ 0.2 small effect size, ≤ 0.5 medium effect size, ≤ 0.8 large effect size; the effect size is large for unoptimized AFS, approximately 3-fold larger for target-optimized AFS, and extremely large for BAFS (∼50-fold increase as compared to unoptimized AFS). Bootstrapping approaches were used to calculate the uncertainty in the metric estimate due to the limited number of runs on each chip. Finally, a non-parametric test was used to ascertain the significance of the separation observed in each method, showing a *P*-value of 0.117 for unoptimized AFS (not statistically significant), 0.029 for target-optimized AFS (statistically significant) and 0.028 for BAFS (statistically significant).

|  | AFS, target: Thp-1 |  | AFS, optimized target: Nalm-6 |  | BAFS, target: NY-ESO-1 9V SCT+ SLB |  |
| --- | --- | --- | --- | --- | --- | --- |
| **Scalar metrics calculated from average binding curves. Variability between runs not taken into account.** Higher values indicate better separation between average binding curves. |  |  |  |  |  |  |
| Area between curves (ΔAUC) | 1.23e+04 | | 2.58e+04 | | 8.18e+04 | |
| MAD | 12.31 |  | 25.76 |  | 81.65 |  |
| RMSD | 12.46 |  | 26.07 |  | 82.39 |  |
| **Effect size metrics (in terms of standard deviation). Incorporate both separation and variability across runs.** Higher effect size indicates better separation and lower variability across runs. Hedges'g applies a correction factor to Cohen's d for small sample sizes (<20). |  |  |  |  |  |  |
| Pooled Cohen's d | 1.26 |  | 3.97 |  | 69.88 |  |
| Hedges' g | 1.09 |  | 3.46 |  | 58.84 |  |
| **Resample runs with replacement, compute metric for each resample, generate distribution. Shows uncertainty in the metric estimate due to limited runs.** |  |  |  |  |  |  |
| Bootstrap ΔAUC CI | 840.31 | 2.39e+04 | 1.84e+04 | 3.37e+04 | 8.03e+04 | 8.34e+04 |
| Bootstrap pooled Cohen's d CI | 0.12 | 4.06 | 3.04 | 12.65 | 64.23 | 2.93e+02 |
| Bootstrap Hedges' g CI | 0.11 | 3.53 | 2.65 | 11.00 | 54.11 | 2.47e+02 |
| Shuffle sample/control labels across runs, recompute metric 5000 times, compute fraction of permutations where difference ≥ observed. **Non-parametric test for significance of separation, accounts for limited number of runs.** | | | | | | |
| Permutation p-value | 0.117 |  | 0.029 |  | 0.028 |  |

## Material and Methods

### Cell lines

All cell lines were maintained at 37°C with 5% CO₂ in RPMI1640 (with phenol red, with stable glutamine, catalog number 392-0427, VWR) supplemented with 10% (V/V) fetal bovine serum (FBS, catalog number A5209502, gibco), 1% (V/V) penicillin/streptomycin (catalog number PS-B, Capricorn Scientific), 1% (V/V) sodium pyruvate (catalog number NPY-B, Capricorn Scientific). Cells were cultured in suspension and passaged every 2-3 days to maintain cell density between 0.2-1 × 10^6^ cells/mL. All cell lines were routinely tested for mycoplasma contamination and maintained under sterile conditions.

### Target cell lines

For CD19-targeting CAR+ Jurkat effector T cells, the target cell of choice was wild-type Nalm-6 cells, a CD19+ precursor B-cell acute lymphoblastic leukemia cell line. For 1G4 αβTCR+ Jurkat effector T cells, the target cell of choice was Thp-1 (human monocytic cell line) or Nalm-6 cells, both expressing HLA-A*02:01 needed for NY-ESO-1 antigen presentation to 1G4 αβTCR+ effector Jurkat T cells.

### Effector cell lines

#### Anti-CD19 CAR expressing cells

For experiments measuring avidity to CD19+ cells, our effector cell of choice consisted of Jurkat e6.1 cells lentivirally transduced to express CAT CAR^21^ and, where applicable, ZAP-70-eGFP. To this end, HEK293T cells were first transfected with the pHR plasmid of interest, psPax and pMD2G using GeneJuice (Sigma-Aldrich, catalog number 70967), following the manufacturer’s protocol. Viral supernatant was collected after 72h and used for Jurkat transduction. CAT CAR+ Jurkat T cells were then sorted for high CAR expression as follows: CAT CAR-transduced Jurkat cells were prepared for sorting first by an incubation with a CD19 detection kit (CD19 CAR Detection Reagent, human, Biotin; catalog number 130-129-550, Miltenyi Biotec) according to manufacturer’s instructions. A second staining was performed with an antibody mix containing streptavidin-PE conjugate antibody (catalog number 36777, Cell Signaling) and pan αβTCR-APC (catalog number IP26, eBioscience) for 30 minutes at 4°C. Cell sorting was performed on the BD FACS Aria II Cell sorter. Jurkat cells with similar αβTCR-APC signal (mean fluorescence intensity (MFI) 1-5 x 10^3^) and high CAR expression (MFI > 1-2 x 10^4^) were collected in 5mL FACS tubes containing 500 µL of RPMI-1640 medium supplemented with 300 U/ml penicillin/streptomycin and 10% fetal calf serum and cultured for the first 2 days at a concentration of 0.5 x 10^6^ cells per mL. 50 IU/mL of interleukin 2 (IL-2, Clinigen) was added to the cells 2 days after sorting to support recovery. More information relevant to CAT CAR+ Jurkat cell sorting is provided in **Supplementary Fig. 1**.

#### CD8ab/1G4αβ TCR expressing cells

All genes were cloned into the second-generation lentiviral backbone pHR under control of the SFFV promoter. Gene identity and integrity was confirmed by Sanger sequencing. TCR α- and β-chains were co-expressed using a P2A self-cleaving sequence. CD8 wild-type and mutant constructs were cloned and verified in the same manner. CD8 wild-type and mutant sequences are provided in **Supplementary Fig. 2**.

Lentivirus was produced in Lenti-X 293T cells (Takara Bio) by co-transfection of pHR transfer plasmid with second-generation packaging plasmids using Fugene 6. Viral supernatant was harvested 48-72 hours post-transfection, clarified by centrifugation, and used fresh or stored at -80°C.

Jurkat cells were transduced with viral supernatant and cultured for 3-5 days before analysis. Expression of transduced TCR and CD8 constructs was confirmed by flow cytometry using fluorophore-conjugated antibodies against TCRαβ, CD3ε and CD8 (**Supplementary Fig. 3** and **4**).

Absolute receptor quantification was performed using Quantibrite PE beads (BD Biosciences, catalog number 340495) according to the manufacturer’s instructions and reported in **Supplementary Fig. 5**. Briefly, PE-conjugated antibodies (BioLegend) were used to stain cells, and fluorescence intensity was converted to molecules per cell using calibration curves generated from Quantibrite beads with a defined number of PE molecules per bead.

### Recombinant single-chain trimer (SCT) production

HLA-A*02:01* single-chain trimers (SCTs) encoding the NY-ESO-1 9V peptide (SLLMWITQV) were generated by genetically linking the peptide, β2-microglobulin, and the HLA-A*\**02:01 heavy chain via flexible glycine-serine linkers. A Y84A mutation was incorporated into the HLA heavy chain to permit linker accommodation within the F pocket.

Recombinant SCT proteins were expressed in Expi293F suspension cells cultured in Expi293 expression medium at 37°C, 5% CO₂, with shaking at 130 rpm. Cells were transfected at a density of 3 × 10⁶ cells/mL using linear polyethyleneimine (PEI) at a 3:1 PEI:DNA ratio. Transfected cultures were maintained for 5-6 days before harvesting supernatant by centrifugation.

SCT proteins were purified by nickel affinity chromatography (Ni-NTA) after overnight incubation with nickel-agarose beads, eluted with 200 mM imidazole followed by size exclusion chromatography (SEC) in HEPES-buffered saline to isolate monomeric protein. Purity and integrity were confirmed by SDS-PAGE under reducing and non-reducing conditions. Proteins were concentrated using centrifugal filters, aliquoted, flash-frozen, and stored at -80°C until use.

### Liposome preparation

For all experiments performed on lipid bilayers containing nickelated lipids, small unilamellar vesicles (SUVs) used for SLB formation were prepared as follows: first, lipids dissolved in chloroform were transferred to a round-bottomed glass flask using precision syringes. The molar ratio of lipids used was 98.5% 1,2-dioleoyl-sn-glycero-3-phosphocholine (DOPC; Avanti Lipids, Alabaster, Alabama, USA), 1% 1,2-dioleoyl-sn-glycero-3-[(N-(5-amino-1-carboxypentyl)iminodiacetic acid) succinyl] (nickel salt)(18:1 DGS-NTA(Ni); Avanti Lipids), and 0.5% ATTO390 conjugated to 1,2-dioleoyl-sn-glycero-3-phosphoethanolamine (DOPE)(ATTO-TEC GmbH, Siegen, Germany). After thorough mixing, the chloroform solution of the lipid mixture was slowly dried under a stream of nitrogen gas while rotating the flask to obtain a thin film of lipids covering the walls of the round-bottomed glass flask. The flask was left to desiccate under vacuum overnight to remove traces of chloroform. The appropriate volume of HEPES buffered saline (HBS; 20 mM HEPES (catalog number H3375-250G, Sigma-Aldrich), 140 mM sodium chloride (catalog number S3014-1KG, Sigma-Aldrich), in milliQ, pH=7.45) warmed to 60°C was then added to the flask slowly, while concomitantly agitating the solution to ensure efficient formation of a 2 mg/mL liposome suspension from the dry lipid film. The multilamellar liposome suspension was then harvested, aliquoted and plunge-frozen in liquid nitrogen for long-term storage. For every experimental day, a tube containing 50 μL of liposome suspension was thawed and ultrasonicated to break up multilamellar vesicles and homogenize liposome size, yielding a translucent SUV suspension. This was further diluted 10-fold with HBS before injecting the liposome suspension into the AFS chips to form SLBs. For ligand density titration assays, where the percentage of 18:1 DGS-NTA(Ni) was titrated down over two orders of magnitude from 1% to 0.01%, 1% nickelated liposomes were mixed either 1:10 (for 0.1% nickelated liposomes) or 1:100 (for 0.01% nickelated liposomes) with liposomes containing 100% DOPC. The mixture was then ultrasonicated prior to use, diluted 10-fold with HBS and injected into the AFS chip for SLB formation (see *Target SLB formation* subsection below). To prepare biotinylated liposomes, a lipid mixture containing 1% (mol) 1,2-dipalmitoyl-sn-glycero-3-phosphoethanolamine-N-(cap biotinyl) (sodium salt) (Biotin-Cap-PE; Cat #870277; Avanti Lipids, Alabaster, Alabama USA), 98.5% DOPC, and 0.5% ATTO390-DOPE was used, with the rest of the liposome preparation protocol identical to that as described above for nickelated liposomes.

### Bilayer fluidity assessment – fluorescence recovery after photobleaching in TIRF

A bilayer composed of DOPC : DGS NTA(Ni) : ATTO 390 DOPE at a 98.5 : 1 : 0.5 molar ratio was formed by incubating SUVs in a z-Movi flow chamber (extracted from a z-Movi chip) for 30 min (Fig. 1a, step 1). After washing out excess liposomes with HBS, the bilayer was first passivated using a 5% solution of casein (catalog number C3400-500G, Sigma-Aldrich) in HBS. 10xHis-tagged CD19 ECD (catalog number CD9-H52H2-500ug, ACROBiosystems) conjugated with Janelia Fluor 646 via random lysine labeling (Janelia Fluor 646, NHS ester, catalog number 6148, Tocris) was suspended in 1% casein solution in HBS to a final concentration of 0.3 μM, and 100 μL flushed into the flow chamber. After 1 hour of incubation, excess CD19 was washed out by pulling through 800 μL of 1% casein solution in HBS. The z-Movi flow chamber was then mounted on a custom built TIRF^22^ microscope and a circular area of the bilayer was photobleached. The diffusion of fluorescent lipid species as well as fluorescent CD19 into the bleached area was tracked over approximately 2 minutes. The mean fluorescence intensity was subsequently double normalized to a reference area and adjusted for photobleaching during post-FRAP acquisition.

### z-Movi chip cleaning and functionalization

LUMICKS z-Movi chips were cleaned using the following procedure: 2×400 μL milliQ water were pulled through the chip, followed by pulling through 3×400 μL 2% Hellmanex III (catalog number 9-307-011-4-507, Hellma GmbH & Co. KG) alkaline cleaning solution in milliQ. The chips were incubated with alkaline cleaning solution overnight and 5×400 μL milliQ was pulled through the chip afterwards. The chips were stored dry prior to use. Before using z-Movi chips, 2×400 μL milliQ was pulled through, followed by pulling in 200 μL 1M NaOH. After 10 minutes, NaOH was washed out from the chip using 3×400 μL milliQ. At this point, the protocol for chip functionalization diverges based on the intended use, as discussed below.

### Target cell monolayer formation

In the case where a cell monolayer was formed as a target for avidity measurements, the chip was incubated with 100 μL 0.01% poly-L-lysine (PLL, catalog number P4707, Sigma-Aldrich) for 1 hour, after which the excess PLL was washed out by pulling 2×400 μL phosphate buffered saline (PBS; catalog number P4417-50TAB, Sigma-Aldrich) through the z-Movi AFS chip. A small volume of serum-free cell medium (RPMI1640, 1% sodium pyruvate, 25 mM HEPES buffer (catalog number BEBP17-737E, Lonza)) was flushed into the chip for storage until target monolayer formation. Target cells were harvested from culture, centrifuged at 300 x g, the pellet was resuspended with 9 mL PBS and cell suspension pelleted at 300 x g. The pellet was resuspended in serum-free cell medium (final cell density ∼1-2 x 10^8^ cells/mL) and flushed into the PLL-coated z-Movi chip. After incubating the cells for 30 minutes, excess target cells were removed by flushing the chip with 3×200 μL serum-free cell medium. Avidity measurements were performed successively with short incubation times, thus no medium refresh was necessary in most cases; if the time before avidity measurements exceeded 2 hours, a medium refresh with complete cell medium (containing 10% FBS) is recommended to maintain viability of target cell monolayer.

### Target SLB formation

In the case where a supported lipid bilayer was used as a target for avidity measurements, 100 μL of 0.2 mg/mL SUVs (for composition, see *Liposome preparation* subsection above) was flushed into the chip immediately after removing NaOH and pulling 400 μL milliQ water through the chip (Fig. 1a, step 1). The SUV suspension was incubated for a minimum of 30 minutes in the dark to prevent fluorescent lipid species bleaching. Excess liposomes were flushed out using 400 μL of HBS. For supported lipid bilayers containing nickelated lipids and functionalized with CD19, a 0.5% casein (w/V) solution in HBS was then flushed into the chip to passivate any glass surfaces in the flow chamber that have not been coated with an SLB to prevent non-specific protein binding. After 30 minutes of passivating the bilayer with 0.5% casein in HBS, the chips was washed by pulling in 400 uL of 0.1% casein in HBS. Then, 100 μL of 0.314 μM solution of His-tagged CD19 ECD (Human CD19 (20-291), His Tag; AcroBiosystems, catalog number, CD9-H52H2-50ug, AcroBiosystems) in 0.1% casein HBS was flushed into the AFS chip and incubated for 1 hour to allow His-tagged CD19 ECD to bind to the nickelated lipids of the SLB (Fig. 1a, step 2). Excess CD19 ECD was removed by pulling 2×400 μL of imaging medium through the microfluidic chip immediately before effector cell sample application. For supported lipid bilayers containing nickelated lipids and functionalized with NY-ESO-1 9V single-chain trimer, bilayer-coated chips were passivated using 100 uL of 5% (w/V) bovine serum albumin (Sigma-Aldrich, catalog number A7906) in HBS for 30 minutes before incubating the bilayer with 100 nM NY-ESO-1 9V SCT in 1% BSA-containing HBS for 1 hour.

For avidity measurements on biotinylated bilayers, 100 uL of HBS containing 1μg/mL NeutrAvidin (Thermo Scientific, catalog number 31000) and 0.1% (w/V) casein was pulled into the z-Movi chip containing passivated SLBs and incubated for 1 hour at room temperature. Excess NeutrAvidin was removed from the chip by pulling through 400 uL HBS. Finally, 100 μL of HBS containing 0.314 μM biotinylated human CD19 ((20-291), His, Avitag; ACROBiosystems, catalog number CD9-H82E9) and 0.1% casein was flushed into the AFS chip and incubated for 1 hour. Excess CD19 ECD was removed by pulling 2×400 uL of imaging medium through the microfluidic chip immediately prior to effector cell sample application.

### Effector cell staining for avidity measurements

Effector cells were harvested during the log phase of cell growth, at ∼10^6^ cells/mL culture density. The number of effector cells used per avidity measurement was ∼2×10^5^. A staining solution consisting of 1.5 μL CellTrace Far Red proliferation dye (catalog number C34564, Invitrogen/Thermo Fisher Scientific) in dimethyl sulfoxide (DMSO, anhydrous, catalog number D12345, Invitrogen) dissolved in 1 mL PBS was prepared. After harvesting, effector cell culture was diluted 1:1 in PBS and centrifuged at 300 x g for 5 minutes. The resulting supernatant was removed and the pellet resuspended in 9 mL PBS and centrifuged at 300 x g. The pellet was then resuspended in 1 mL staining solution per 2 million cells, followed by 15 minutes incubation in dark at 37°C. Afterwards, imaging medium (RPMI1640 (without phenol red, without L-glutamine, catalog number 392-0430, VWR), 10% FBS, 1% GlutaMAX (catalog number 35050-038, gibco), 20 mM HEPES buffer) was added at a 1:5 (V/V) ratio (staining solution : imaging medium) and incubated in the dark for 5 minutes at 37°C to quench the staining reaction. The stained cell suspension was centrifuged at 300 x g, after which the supernatant was aspirated and the pellet was resuspended in imaging medium to a final working concentration of ∼10^7^ cells/mL, with the individual sample volume being 20 μL (∼2 x 10^5^ cells). The working suspension of effector cells was kept in a 96-well plate in a cell incubator prior to use.

### Avidity measurements

After functionalizing the z-Movi AFS microfluidic chip with either target cell monolayers or supported lipid bilayers as described above, avidity measurements were performed using the Oceon software for the z-Movi setup (version 1.5.5), as provided by LUMICKS. For each chip, 20 μL of effector cell sample were slowly pulled into the microfluidic chip using a 3 mL syringe (Fig. 1a, step 3). The sample reservoir was then rinsed 3 times with 400 μL of imaging medium, leaving 200 μL of imaging medium in the reservoir. After 5 minutes of incubation (Fig. 1a, step 4), a force ramp was applied, from 0 to 1000 pN in 1.333 pN increments (Fig. 1a, step 5). Cell tracking and counting was performed on the fly using the proprietary Oceon software. After each avidity run, the detached effector cells were flushed out from the chip using imaging buffer. For consecutive avidity measurements (as described in Extended Data Fig. 2a, bottom), the procedure described above was repeated up to 6 times, whereas for parallel avidity measurements on different chips (as described in Extended Data Fig. 2a, top) each chip was removed from the setup after an experimental run and replaced with a new chip for replicate measurements. After use, microfluidic AFS chips were cleaned using the approach described above.

### Data analysis and visualization

We analyzed our data within a survival analysis framework, in which applied force was treated as a time-like variable and cell unbinding events were treated as failures. Depending on condition, a substantial fraction of cells remained bound at the maximum applied force (1000 pN) in our (B)AFS data, resulting in right-censored measurements. Each observation consisted of a force value and a binary indicator denoting event occurrence (unbinding) or right-censoring (cells remaining bound at the maximum applied force of 1000 pN). Data from individual experiments were pooled by experimental condition. Each condition comprised n ≥ 2 replicates.

Kaplan–Meier survival curves were computed using a custom MATLAB implementation, with survival defined as the probability of a cell remaining bound at a given force. Confidence intervals were estimated using Greenwood’s formula and visualized alongside survival curves. Pairwise comparisons between conditions were performed using the log-rank test, implemented in a custom function that computes the Mantel–Haenszel statistic and corresponding hazard ratios. *P*-values were adjusted for multiple comparisons using Bonferroni correction. Effect sizes were quantified using hazard ratios, representing the relative instantaneous rate of unbinding between conditions. To identify practically significant differences, comparisons were considered significant only if they satisfied both a statistical threshold (Bonferroni-adjusted *P*-value < 0.05) and an effect size criterion (hazard ratio (HR) ≥ 2 or ≤ 0.5).

Raw data (cell unbinding force and average binding curves) were extracted using Oceon (version 1.5.5). Survival analysis and data visualization was performed using custom Matlab (version R2024b, MathWorks) code, based on kmplot (version 2.0.0.0) and logrank (version 2.0.0.0) functions available through Matlab File Exchange / GitHub (Giuseppe Cardillo (2026). KMplot (https://github.com/dnafinder/kmplot), GitHub. Retrieved March 22, 2026. Logrank (https://github.com/dnafinder/logrank), GitHub. Retrieved March 22, 2026).

## Acknowledgements

We thank Marcel Winter (AMOLF) for help with maintaining the cell lines. We thank Lucrezia Gatti (University Medical Center Utrecht) for help with sorting cells. We acknowledge support from AMOLF technical engineering departments. We thank LUMICKS BV (Amsterdam, Netherlands) for technical support and for constructive feedback on the manuscript. N.J. is supported by EMBO (ALTF 896-2022) and HFSP (LT0031/2023-L1) postdoctoral fellowships. T.M.J.E. was funded by project 24PPS068 (CARTTECH) of the PPS-Programmatoeslag of Holland High Tech. C.N. was funded by project number VI.Vidi.203.037 of the Vidi Talent Programme funded by the Dutch Research Council (NWO). K.A.G. also acknowledges support by the WISE program of NWO and Oncode. M.V., A.C.H. and R.A.F. are supported by the Chinese Academy of Medical Sciences (CAMS) Innovation Fund for Medical Science (CIFMS), China (grant number: 2024-I2M-2-001-1) and Cancer Research UK (DRCCIP-Nov23/100004). A.C.H. is supported by a Wellcome Early-Career Award (227585/Z/23/Z)

## Contributions

N.J., S.J.T. and K.A.G. conceived the study. N.J. and T.M.J.E. designed the experiments and wrote the original draft of the manuscript. N.J., T.M.J.E. and A.W. performed avidity measurements and data analysis. C.N. contributed to the development of the methodology and provided resources (CAT CAR lentiviral transduction and cell sorting). M.V., A.C.H. and R.A.F. provided resources (TCR/CD8 lentiviral transduction and cell sorting, NY-ESO 1 9V peptide and HLA-A02:01 NY-ESO-1 9V SCT samples). R.A.F. conceived TCR/CD8 experiments. N.J., S.J.T. and K.A.G. wrote the final version of the paper. S.J.T. and K.A.G. supervised the project. All authors commented on and approved the final version of the manuscript.

## Ethics declarations

## Competing interests

The authors declare no competing interests.

## Notes

### Competing Interest Statement

The authors have declared no competing interest.

## References

1. Baulu, E., Gardet, C., Chuvin, N. & Depil, S. TCR-engineered T cell therapy in solid tumors: State of the art and perspectives. Sci. Adv. 9, eadf3700 (2023).

2. Zugasti, I. et al. CAR-T cell therapy for cancer: current challenges and future directions. Sig Transduct Target Ther 10, 210 (2025).

3. Rosenberg, S. A. & Restifo, N. P. Adoptive cell transfer as personalized immunotherapy for human cancer. Science 348, 62–68 (2015).

4. Zhang, Y. et al. Safety and efficacy of a novel anti-CD19 chimeric antigen receptor T cell product targeting a membrane-proximal domain of CD19 with fast on- and off-rates against non-Hodgkin lymphoma: a first-in-human study. Mol Cancer 22, 200 (2023).

5. Xiong, Y., Libby, K. A. & Su, X. The physical landscape of CAR-T synapse. Biophysical Journal 123, 2199–2210 (2024).

6. Considerations for the Development of Chimeric Antigen Receptor (CAR) T Cell Products - Guidance for Industry.

7. Kamsma, D. et al. Single-Cell Acoustic Force Spectroscopy: Resolving Kinetics and Strength of T Cell Adhesion to Fibronectin. Cell Reports 24, 3008–3016 (2018).

8. Kamsma, D., Creyghton, R., Sitters, G., Wuite, G. J. L. & Peterman, E. J. G. Tuning the Music: Acoustic Force Spectroscopy (AFS) 2.0. Methods 105, 26–33 (2016).

9. Barisa, M. et al. Functional avidity of anti-B7H3 CAR-T constructs predicts antigen density thresholds for triggering effector function. Nat Commun 16, 7196 (2025).

10. Halim, L. et al. Engineering of an Avidity-Optimized CD19-Specific Parallel Chimeric Antigen Receptor That Delivers Dual CD28 and 4-1BB Co-Stimulation. Front. Immunol. 13, 836549 (2022).

11. Larson, R. C. et al. CAR T cell killing requires the IFNγR pathway in solid but not liquid tumours. Nature 604, 563–570 (2022).

12. Davis, S. J. & Van Der Merwe, P. A. The structure and ligand interactions of CD2: implications for T-cell function. Immunology Today 17, 177–187 (1996).

13. Davis, S. J. et al. The nature of molecular recognition by T cells. Nat Immunol 4, 217–224 (2003).

14. Shilts, J. & Wright, G. J. Mapping the Human Cell Surface Interactome: A Key to Decode Cell-to-Cell Communication. Annual Review of Biomedical Data Science 7, 155–177 (2024).

15. Shilts, J. et al. A physical wiring diagram for the human immune system. Nature 608, 397–404 (2022).

16. Hein, M. Y. et al. A Human Interactome in Three Quantitative Dimensions Organized by Stoichiometries and Abundances. Cell 163, 712–723 (2015).

17. Wojtowicz, W. M. et al. A Human IgSF Cell-Surface Interactome Reveals a Complex Network of Protein-Protein Interactions. Cell 182, 1027–1043.e17 (2020).

18. Gérard, A., Cope, A. P., Kemper, C., Alon, R. & Köchl, R. LFA-1 in T cell priming, differentiation, and effector functions. Trends in Immunology 42, 706–722 (2021).

19. Bertolet, G. & Liu, D. The Planar Lipid Bilayer System Serves as a Reductionist Approach for Studying NK Cell Immunological Synapses and Their Functions. in Natural Killer Cells (ed. Somanchi, S. S.) vol. 1441 151–165 (Springer New York, New York, NY, 2016).

20. Dustin, M. L. Insights into Function of the Immunological Synapse from Studies with Supported Planar Bilayers. in Immunological Synapse (eds Saito, T. & Batista, F. D.) vol. 340 1–24 (Springer Berlin Heidelberg, Berlin, Heidelberg, 2010).

21. Ghorashian, S. et al. Enhanced CAR T cell expansion and prolonged persistence in pediatric patients with ALL treated with a low-affinity CD19 CAR. Nat Med 25, 1408–1414 (2019).

22. Niederauer, C., Seynen, M., Zomerdijk, J., Kamp, M. & Ganzinger, K. A. The K2: Open-source simultaneous triple-color TIRF microscope for live-cell and single-molecule imaging. HardwareX 13, e00404 (2023).

23. Hsu, C.-J. et al. Ligand Mobility Modulates Immunological Synapse Formation and T Cell Activation. PLoS ONE 7, e32398 (2012).

24. Wang, Y. et al. Protocol to measure cell avidity between cord blood-derived NK cells and leukemia cell line KG-1a. STAR Protocols 5, 103387 (2024).

25. Dam, T., Junghans, V., Humphrey, J., Chouliara, M. & Jönsson, P. Calcium Signaling in T Cells Is Induced by Binding to Nickel-Chelating Lipids in Supported Lipid Bilayers. Front. Physiol. 11, 613367 (2021).

26. Chen, Y.-T. et al. A testicular antigen aberrantly expressed in human cancers detected by autologous antibody screening. Proc. Natl. Acad. Sci. U.S.A. 94, 1914–1918 (1997).

27. Gao, G. F. et al. Crystal structure of the complex between human CD8αα and HLA-A2. Nature 387, 630–634 (1997).

28. Kern, P. S. et al. Structural Basis of CD8 Coreceptor Function Revealed by Crystallographic Analysis of a Murine CD8αα Ectodomain Fragment in Complex with H-2Kb. Immunity 9, 519–530 (1998).

29. Garcia, K. C. et al. CD8 enhances formation of stable T-cell receptor/MHC class I molecule complexes. Nature 384, 577–581 (1996).

30. Jiang, N. et al. Two-Stage Cooperative T Cell Receptor-Peptide Major Histocompatibility Complex-CD8 Trimolecular Interactions Amplify Antigen Discrimination. Immunity 34, 13–23 (2011).

31. Xu, X.-N. et al. A Novel Approach to Antigen-Specific Deletion of CTL with Minimal Cellular Activation Using α3 Domain Mutants of MHC Class I/Peptide Complex. Immunity 14, 591–602 (2001).

32. Artyomov, M. N., Lis, M., Devadas, S., Davis, M. M. & Chakraborty, A. K. CD4 and CD8 binding to MHC molecules primarily acts to enhance Lck delivery. Proc. Natl. Acad. Sci. U.S.A. 107, 16916–16921 (2010).

33. Van Der Merwe, P. A. & Cordoba, S.-P. Late Arrival: Recruiting Coreceptors to the T Cell Receptor Complex. Immunity 34, 1–3 (2011).

34. Xu, H. & Littman, D. R. A kinase-independent function of Lck in potentiating antigen-specific T cell activation. Cell 74, 633–643 (1993).

35. Malissen, B. & Bongrand, P. Early T Cell Activation: Integrating Biochemical, Structural, and Biophysical Cues. Annu. Rev. Immunol. 33, 539–561 (2015).

36. Kim, P. W., Sun, Z.-Y. J., Blacklow, S. C., Wagner, G. & Eck, M. J. A Zinc Clasp Structure Tethers Lck to T Cell Coreceptors CD4 and CD8. Science 301, 1725–1728 (2003).

37. Srinivasan, S., Zhu, C. & McShan, A. C. Structure, function, and immunomodulation of the CD8 co-receptor. Front. Immunol. 15, 1412513 (2024).

38. Wyer, J. R. et al. T Cell Receptor and Coreceptor CD8αα Bind Peptide-MHC Independently and with Distinct Kinetics. Immunity 10, 219–225 (1999).

39. Pettmann, J. et al. The discriminatory power of the T cell receptor. eLife 10, e67092 (2021).

40. Bachmann, M. F. et al. Distinct Roles for LFA-1 and CD28 during Activation of Naive T Cells: Adhesion versus Costimulation. Immunity 7, 549–557 (1997).

41. Salzer, B. et al. Engineering AvidCARs for combinatorial antigen recognition and reversible control of CAR function. Nat Commun 11, 4166 (2020).

42. Albayrak, G., Wan, P. K.-T., Fisher, K. & Seymour, L. W. T cell engagers: expanding horizons in oncology and beyond. Br J Cancer 133, 1241–1249 (2025).

43. Majzner, R. G. & Mackall, C. L. Tumor Antigen Escape from CAR T-cell Therapy. Cancer Discovery 8, 1219–1226 (2018).

44. Caruso, H. G. et al. Tuning Sensitivity of CAR to EGFR Density Limits Recognition of Normal Tissue While Maintaining Potent Antitumor Activity. Cancer Research 75, 3505–3518 (2015).

45. Céspedes, P. F., Beckers, D., Dustin, M. L. & Sezgin, E. Model membrane systems to reconstitute immune cell signaling. The FEBS Journal 288, 1070–1090 (2021).

46. Hwang, L. Y., Götz, H., Knoll, W., Hawker, C. J. & Frank, C. W. Preparation and Characterization of Glycoacrylate-Based Polymer-Tethered Lipid Bilayers on Benzophenone-Modified Substrates. Langmuir 24, 14088–14098 (2008).

