## Supplementary Information for "Bilayer acoustic force spectroscopy (BAFS) for quantifying receptor-antigen binding strength in immune synapses"

**Supplementary Figure 1.**

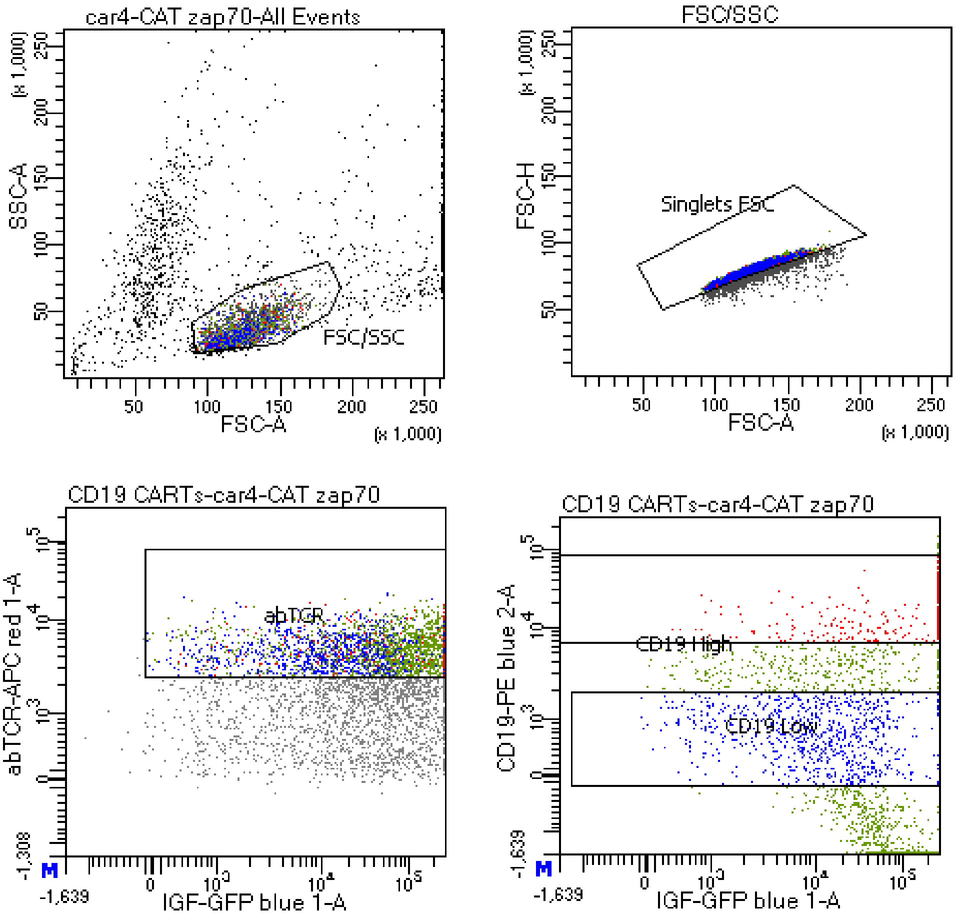


| Population | #Events | %Parent | CD19-PE b... Mean | abTCR-AP... Mean |
|---|---|---|---|---|
| CD19 High | 259 | 9.9 | 17,124 | 5,351 |
| abTCR | 2,605 | 57.7 | 1,052 | 5,054 |
| CD19 Low | #### | #### | 748 | 4,957 |

| Population | #Events | %Parent | %Total |
|---|---|---|---|
| All Events | 10,000 | #### | 100.0 |
| FSC/SSC | 6,226 | 62.3 | 62.3 |
| Singlets FSC | 4,513 | 72.5 | 45.1 |
| abTCR | 2,605 | 57.7 | 26.0 |
| CD19 High | 259 | 9.9 | 2.6 |
| CD19 Low | 820 | 31.5 | 8.2 |

**Supplementary Fig. 1: Cell sorting information for Jurkat e6.1 cells expressing CAT CAR and ZAP70-GFP.** Sorting was performed as follows: CAT CAR-transduced Jurkat cells were prepared for sorting by a first incubation with CD19 detection kit (CD19 CAR Detection Reagent, human, Biotin; catalog number 130-129-550, Miltenyi Biotec) according to manufacturer's instructions. A second staining was performed with an antibody mix containing streptavidin-PE conjugate antibody (catalog number 36777, Cell Signaling) and pan αβTCR-APC (catalog number IP26, eBioscience) for 30 minutes at 4°C. Cells were first gated for live single cells, followed by gating for cells expressing similar levels of αβTCR expression (MFI 1-5 x $10^3$), and lastly for high (CD19-PE high; MFI > 1-2x $10^4$) or low (CD19-PE low; MFI < 1 x$10^3$) CD19 expression. Cell sorting was performed on the BD FACSAria II Cell sorter. Jurkat cells with comparable αβTCR-APC signal (mean fluorescence intensity (MFI) 1.5 x $10^4$) were collected.

**Supplementary Figure 2.**

pHR-hCD8α

SQFRVSPLDRTWNLGETVELKCQVLLSNPTSGCSWLFQPRGAAASPTFLLYLSQNKPKAAEGLD
TQRFSGKRLGDTFVLTLSDFRRENEGYYFCSALSNSIMYFSHFVPVFLPAKPTTTPAPRPPTPAPT
IASQPLSLRPEACRPAAGGAVHTRGLDFACDIYIWAPLAGTCGVLLLSLVITLYCNHRNRRRVCKC
PRPVVKSGDKPSLSARYV

pHR-hCD8α$^{C>S}$

SQFRVSPLDRTWNLGETVELKCQVLLSNPTSGCSWLFQPRGAAASPTFLLYLSQNKPKAAEGLD
TQRFSGKRLGDTFVLTLSDFRRENEGYYFCSALSNSIMYFSHFVPVFLPAKPTTTPAPRPPTPAPT
IASQPLSLRPEACRPAAGGAVHTRGLDFACDIYIWAPLAGTCGVLLLSLVITLYCNHRNRRRVSKS
PRPVVKSGDKPSLSARYV

pHR-hCD8β

LQQTPAYIKVQTNKMVMLSCEAKISLSNMRIYWLRQRQAPSSDSHHEFLALWDSAKGTIHGEEVE
QEKIAVFRDASRFILNLTSVKPEDSGIYFCMIVGSPELTFGKGTQLSVVDFLPTTAQPTKKSTLKKR
VCRLPRPETQKGPLCSPITLGLLVAGVLVLLVSLGVAIHLCCRRRRARLRFMKQFYK

**Supplementary Fig. 2: Sequences of CD8α$_{wt}$, CD8α$_{mut,}$ and CD8β$_{wt}$.** Exchange of cysteine residues C194 and C196 for serine in CD8α disrupts recruitment of Lck to the cytosolic domains of CD8 but has no effect on the binding of CD8 to MHC via its extracellular domain.

**Supplementary Figure 3.**

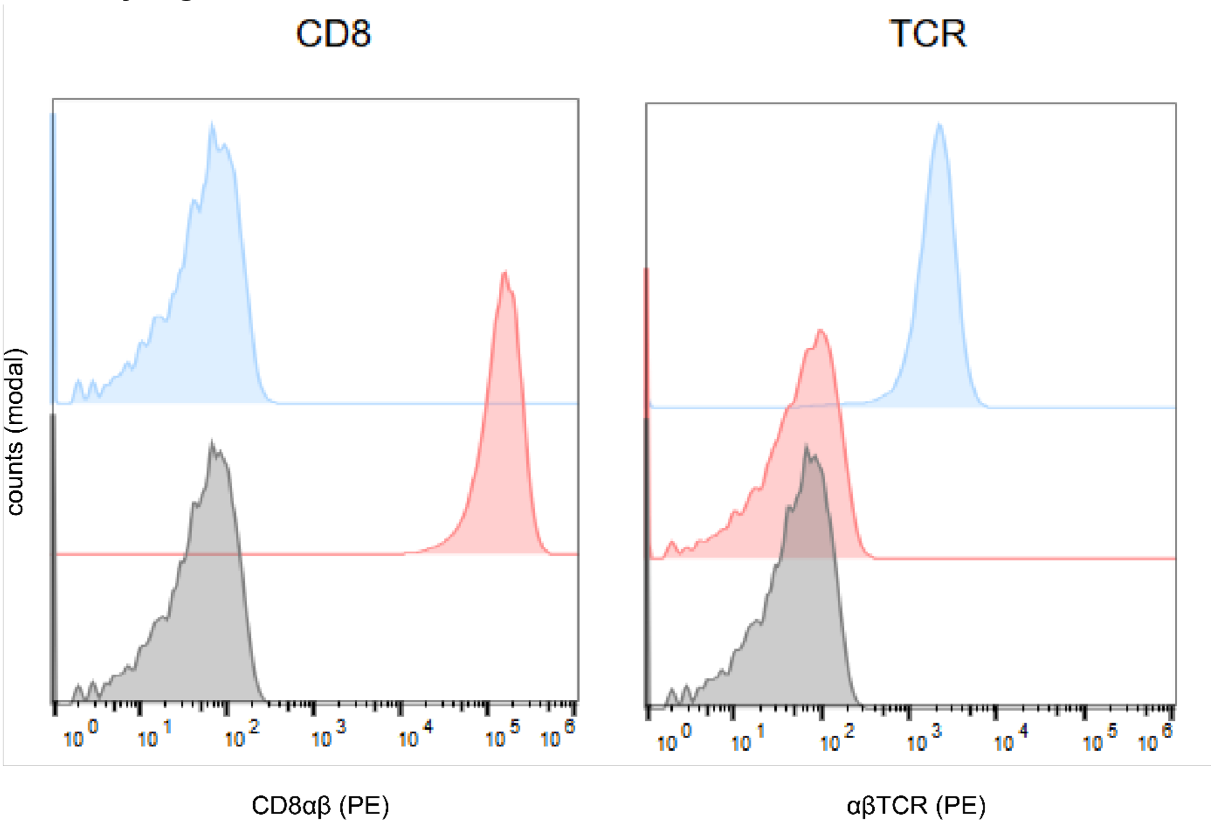


| | Sample Name | Count | Geometric Mean : YL1-A |
|---|---|---|---|
| | D7.fcs | 25693 | 7.79 |
| | D8.fcs | 16744 | 134991 |
| | A2.fcs | 22502 | 8.24 |

| | Sample Name | Count | Geometric Mean : YL1-A |
|---|---|---|---|
| | E7.fcs | 24549 | 1775 |
| | E6.fcs | 15036 | 7.09 |
| | A2.fcs | 22502 | 8.24 |

**Supplementary Fig. 3: Quantification of TCR and CD8 surface expression in Jurkat T cells.** Surface expression levels were assessed by flow cytometry in TCR-deficient Jurkat T cells transduced with lentiviral vectors encoding either the 1G4 αβTCR (blue) or CD8αβ (red). Representative histograms are shown.

**Supplementary Figure 4.**

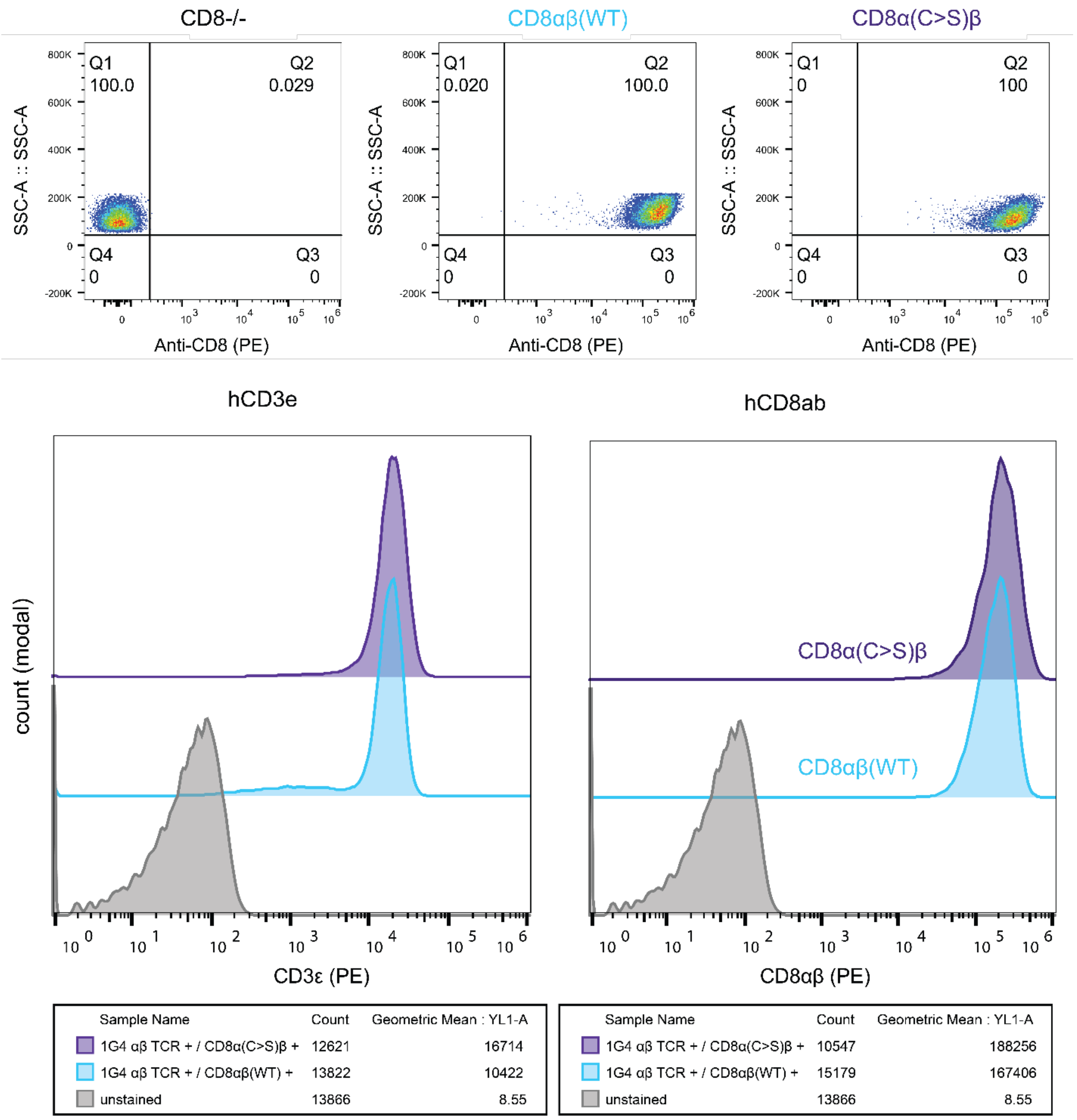


**Supplementary Fig. 4: Quantification of wild-type and CS mutant CD8 surface expression in Jurkat T cells.** Surface expression levels were assessed by flow cytometry in 1G4 TCR-transduced Jurkat T cells following lentiviral transduction with either wild-type CD8αβ (blue) or CS mutant CD8αβ (purple). Representative histograms are shown.

**Supplementary Figure 5.**

| 1G4 TCR | CD8 | # CD8 mol/cell |
|---|---|---|
| - | WT | 737,861 |
| + | WT | 816,029 |
| + | CS | 919,488 |

**Supplementary Fig. 5: Quantification of wild-type and C>S mutant CD8αβ surface expression.** Surface CD8αβ expression (molecules per cell) was determined by flow cytometry following staining with PE-conjugated anti-CD8 antibodies and quantification using QuantiBrite™ calibration beads. Values represent the average number of molecules per cell calculated from >2,000 cells analysed per cell line.

**Supplementary Note 1.**

We tested whether single SLBs can be used for multiple measurements in succession (Extended Data Fig. 1a). For 6 consecutive runs on the same CD19-functionalized SLB chip, we obtained an average of 74.6±6.2% for the percentage of CAR+ T cells that remained bound (Extended Data Fig. 1b). When performing 6 runs in parallel across 6 different chips, the average was 75.2±2.5%, which is in good agreement with the sequential measurements on a single chip. However, data from sequential measurements on a single SLB showed a generally decreasing percentage of cells that remained bound at the endpoint of each successive run, in particular after the fifth consecutive run. As a result, the consecutive data set showed a 3-fold larger coefficient of variation than the data taken in parallel, which was true across the measured force range (Extended Data Fig. 1c). Various effects can contribute to this variation and decrease, including effector cells that remain bound after the experiment and hence decrease the available binding area for the next effector cell sample, washing steps between consecutive runs that may remove CD19 from the SLB, and possibly irreversible removal of CD19 ECD from bilayers by detaching cells. Overall, however, the method allows for consecutive measurements on the same SLB at the cost of slightly increased inter-sample variability.
